# A three-factor model can explain symmetry breaking and the n-1 rule during X-chromosome inactivation

**DOI:** 10.64898/2026.11.03.764019

**Authors:** Kaiyun Guan, Jerry M. Liu, J. Mauro Calabrese, Timothy C. Elston

## Abstract

X-chromosome inactivation (XCI) is a mammalian gene regulatory process that via chromosome-scale transcriptional silencing, leaves active a single X chromosome per diploid genome complement. The mechanisms that enable cells to select the appropriate number of X chromosomes to inactivate while leaving others active remain unclear. We demonstrate that all known chromosomal counting scenarios during XCI can be accounted for by a three-factor mathematical model that includes (1) *Xist*, the long noncoding RNA (lncRNA) responsible for gene silencing during XCI in eutherians, (2) an X-linked transcriptional activator of *Xist*, and (3) the autosomally-encoded SPEN, an *Xist* RNA-binding protein that is essential for gene silencing during XCI. In our model, symmetry between active X chromosomes is broken by stochastic differences in the number of *Xist* molecules that associate with the X chromosomes at the onset of XCI. Subsequently, integrated positive and negative feedback loops amplify then propagate superstoichiometric assemblies of SPEN complexed with *Xist* on the inactive chromosome(s), and select for levels of X-linked activator that enable only one X chromosome per diploid genome complement to remain active. Thus, asymmetric propagation of the SPEN-*Xist* ribonucleoprotein complex may play an essential role in symmetry breaking and chromosomal counting during XCI, independently of or in concert with cis-acting networks of DNA regulatory elements. Our results also have conceptual implications for gene regulatory circuits on autosomes.

**Significance Statement:** X-chromosome Inactivation (XCI) is a mammalian gene regulatory process in which one of two X chromosomes in each female cell is randomly selected for silencing. How cells select only one X for silencing is not understood. We use mathematical modeling to demonstrate that a mechanism in which positive feedback amplifies fluctuations in molecular abundances is sufficient to explain XCI. In particular, the X-linked noncoding RNA *Xist* and the RNA-binding protein SPEN form a positive feedback loop that propagates a ribonucleoprotein complex along the X chromosomes in all-or-none fashion. Our model demonstrates how the asymmetric propagation of an RNA-protein effector complex might establish monoallelic gene expression during XCI. Analogous RNA-driven regulatory circuits may underlie other instances of monoallelic gene expression.

## Introduction

X-chromosome inactivation (XCI) occurs early during the development of all female mammals, and results in the transcriptional silencing of a single X chromosome in every diploid cell. XCI evolved as part of the mechanism to compensate for the dosage imbalance between female mammals, whose genomes harbor two X chromosomes, and male mammals, whose genomes harbor only one. Once chosen for inactivation, the inactive X is propagated in a silenced state essentially for life, marking XCI as one of the most striking known examples of a stable epigenetic switch. Accordingly, since its discovery over 60 years ago, XCI has served as a paradigm of epigenetic regulation from which many aspects of gene regulation have been learned (1).

Critical for the success of XCI is an intrinsic counting mechanism in which cells inactivate a single X chromosome for every diploid genome complement, referred to as n-1 rule, where n represents the total number of X chromosomes per diploid genome equivalent, and “n-1” refers to the number of X chromosomes that become inactivated (2–5). Thus, male cells that harbor only one X chromosome inactivate none, whereas female cells that harbor two X chromosomes inactivate one. Likewise, male and female cells that are otherwise diploid, but harbor supernumerary copies of their X chromosomes inactivate the number of X’s needed such that only one remains active. For example, XXY male cells inactivate one X chromosome, XXX female cells inactivate two, and XXXX tetraploid cells (4n chromosomal content) inactivate two X chromosomes and leave the remaining two X’s active.

The molecule responsible for orchestrating XCI in eutherian mammals is the long noncoding RNA *XIST* (*Xist* in mouse), whose expression from the future inactive X chromosome ultimately leads to stable silencing *in cis*, i.e. along the X chromosome from which *XIST/Xist* is being expressed. At steady state, an inactive X chromosome produces about 100-200 molecules of *XIST/Xist*, which form about 50-100 distinct foci that surround its gene-rich dense regions (6–9). *XIST/Xist* recruits a local assembly of proteins that then coordinate gene silencing during XCI. Among the set of *XIST-*recruited proteins, an RNA-binding protein called SPEN has consistently been shown to be critical during the earliest stages of XCI. SPEN is recruited to a conserved region of *XIST/Xist* and likely works together with a cohort of effector proteins to initiate and then stabilize XCI (7, 10–32).

In pre- and peri-implantation embryos from nearly all eutherians examined to date (including human, monkey, rabbit, and mouse), prior to the developmental timepoint at which full-scale gene silencing from XCI is observed, *XIST/Xist* can exhibit a symmetric expression pattern from both X chromosomes in female and is even expressed on the single X chromosome in males (33–38). The mechanistic basis for the transition from biallelic to monoallelic *XIST/Xist* expression – referred to as “symmetry breaking” – remains unclear. Genetic and computational models suggest that symmetry breaking is governed by a balance of diffusible factors, located on autosomes and X chromosomes, that collectively act through a network of DNA regulatory elements to control the transcription of *XIST/Xist* (39–42). For example, in mouse embryonic stem cells, REX1, an autosomally-encoded protein, represses *Xist* expression by binding to its promoter, while an X-linked protein, RNF12/RLIM, activates *Xist* expression by inducing REX1 degradation (43, 44). Cells lacking REX1 or RLIM can still adhere to the n-1 rule, suggesting that additional factors play important and potentially redundant roles in X chromosomal counting (44–46). To that point, additional X-linked genes, KDM6A, KDM5C, and ZIC3, can activate *Xist* and exhibit properties consistent with roles in symmetry breaking (41, 47–49). These and other proteins, including autosomally-encoded factors, are thought to integrate signals between cis-acting DNA regulatory elements to yield the correct number of inactive X chromosomes upon the completion of XCI (35, 39–41, 49).

We show that all known X chromosomal counting scenarios can be accurately accounted for using a simple three-factor mathematical model. This model is comprised of an X-linked diffusible activator of *Xist* expression, the autosomally-encoded SPEN, and *Xist* itself. In our model, asymmetry first arises due to stochastic differences in the levels of *Xist* expression between its two alleles. Subsequently, asymmetry is amplified in a manner that depends on the concentration of the *Xist* activator, the overall concentration of SPEN, and the propensity of *Xist* to nucleate and propagate stable super-stoichiometric assemblies of SPEN. Thus, through an ability to propagate asymmetrically, the SPEN-*Xist* ribonucleoprotein complex may promote its own monoallelism, independently of or in concert with networks of cis-acting DNA regulatory elements. Analogous mechanisms of RNA-driven protein assembly and positive feedback may likewise promote other instances of monoallelic gene expression.

## Results

### A mathematical model to describe chromosomal counting and symmetry breaking during XCI

Models for random X chromosome inactivation must provide a mechanism for symmetry breaking and enforce the n-1 rule. Symmetry breaking refers to the requirement that X chromosome inactivation occurs in the absence of preexisting landmarks that target specific chromosomes for silencing. The n-1 rule states that if n X chromosomes are present in an otherwise diploid cell, then n-1 of the X chromosomes are transcriptionally silenced. With these constraints in mind, we devised a model for random XCI that consists of 4 elements: 1) positive feedback between *XIST/Xist* and the transcriptional repressor SPEN, 2) a common pool of free SPEN, 3) an X-linked trans-acting transcriptional activator for *Xist* and 4) a negative feedback loop formed by SPEN’s repression of the transcriptional activator (Fig. 1A). Our models focus on SPEN because it has consistently been identified as the single most essential *Xist* co-factor, especially during the earliest stages of XCI (10–19), but we acknowledge that *Xist* also engages SPEN-independent pathways to promote gene silencing, particularly in differentiated cells (24–32). When referring to the *XIST/Xist* gene below, we default to the mouse nomenclature, *Xist*.

**Figure 1.**
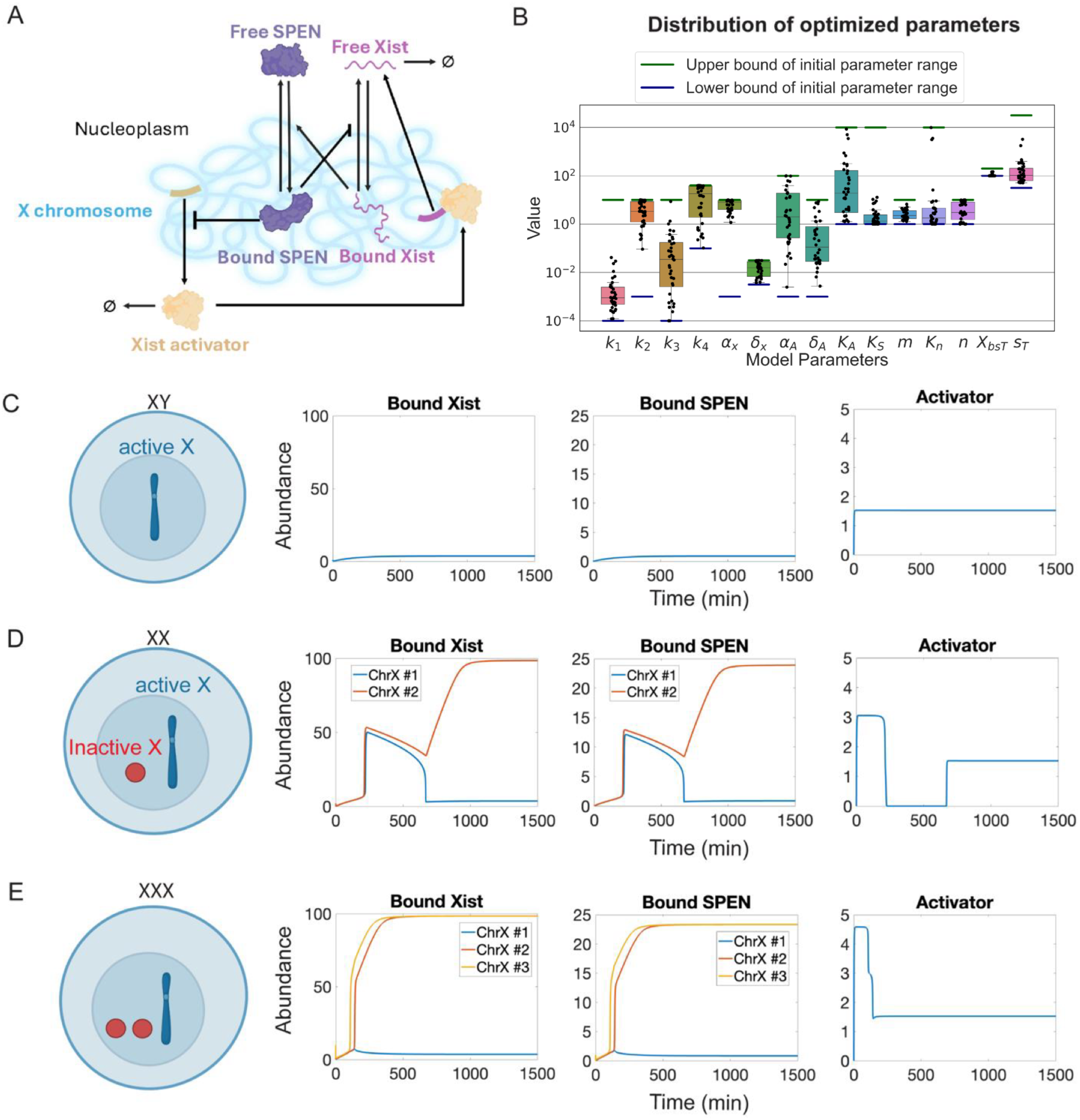
Activator inhibition model for XCI. **A)** Model schematic. Created with BioRender.com. **B)** Parameter sets identified by the evolutionary algorithm. Each dot represents one parameter set (N=36). See Table 1 for parameter descriptions and relevant references. **C)** Time series for number of molecules of bound *Xist*, bound SPEN, and activator (i.e. Abundance) for the case of XY males. The total amount of SPEN molecules per cell in each simulation is S_T_ = 1000. **D)** Same as **C** except for XX females. **E)** Same as **C** except for XXX females.

**Table 1.** Summary of ODE Model Parameters.

| Parameter description | Symbol | Search range | Parameter values used in Fig. 1 and Fig. 3 | Parameter values used in Fig. 2 | Parameter values used in Fig. 4 | Reference |
|---|---|---|---|---|---|---|
| *Xist*-chromosome association rate | *k*<sub>1</sub> | [10<sup>-4</sup>, 10] min<sup>-1</sup> | 0.0084 min<sup>-1</sup> | 0.002 min<sup>-1</sup> | 0.0024 min<sup>-1</sup> |  |
| *Xist*-chromosome dissociation rate in the absence of SPEN | *k*<sub>2</sub> | [10<sup>-3</sup>,10] min<sup>-1</sup> | 8.7 min<sup>-1</sup> | 4.54 min<sup>-1</sup> | 4.21 min<sup>-1</sup> | (7) |
| SPEN-*Xist* association rate | *k*<sub>3</sub> | [10<sup>-4</sup>,10] min<sup>-1</sup> | 0.00026 min<sup>-1</sup> | 8.26 min<sup>-1</sup> | 0.014 min<sup>-1</sup> |  |
| SPEN-*Xist* dissociation rate | *k*<sub>4</sub> | [10<sup>-1</sup>,10<sup>1.6</sup>] min<sup>-1</sup> | 10.2 min<sup>-1</sup> | 0.11 min<sup>-1</sup> | 8.78 min<sup>-1</sup> | (7) |
| Maximum *Xist* transcription rate | *α*<sub>*x*</sub> | [10<sup>-3</sup>,10] min<sup>-1</sup> | 3.2 min<sup>-1</sup> | 9.0 min<sup>-1</sup> | 5.9 min<sup>-1</sup> | (84) |
| Degradation rate for free *Xist* | *δ*<sub>*x*</sub> | [10<sup>-2.5</sup>,10<sup>-1.5</sup>] min<sup>-1</sup> | 0.0076 min<sup>-1</sup> | 0.031 min<sup>-1</sup> | 0.0041 min<sup>-1</sup> | (10, 51, 84–86) |
| Maximum activator production rate | *α*<sub>*A*</sub> | [10<sup>-3</sup>,10<sup>2</sup>] min<sup>-1</sup> | 1.1 min<sup>-1</sup> | 12.8 min<sup>-1</sup> | 66.8 min<sup>-1</sup> |  |
| Degradation rate for the activator | *δ*<sub>*A*</sub> | [10<sup>-3</sup>,10] min<sup>-1</sup> | 0.72 min<sup>-1</sup> | 0.13 min<sup>-1</sup> | 0.048 min<sup>-1</sup> |  |
| Number of activator molecules at which the *Xist* transcription rate is half max | *K*<sub>*A*</sub> | [1,10<sup>4</sup>] | 16.1 molecules | 374.6 molecules | 76.5 molecules | (87–90) |
| Number of SPEN molecules at which the *Xist* dissociation rate is half max | *K*<sub>*S*</sub> | [1,10<sup>4</sup>] | 2.06 molecules | 4.3 molecules | 7.4 molecules | (87–90) |
| Hill coefficient for SPEN inhibition of *Xist* dissociation | m | [1,10] | 3.09 | 2.0 | 3.85 |  |
| Number of SPEN molecules at which the activator production rate is half max | *K*<sub>*n*</sub> | [1,10<sup>4</sup>] | 2.2 molecules | 7.13 molecules | 1.0 | (87–90) |
| Hill coefficient for SPEN inhibition of activator production | n | [1,10] | 9.4 | 1.0 | 10.0 |  |
| Total *Xist* binding sites per chromosome | *X*<sub>*bs*</sub> | [10<sup>2</sup>,10<sup>2.3</sup>] | 100 molecules | 100 molecules | 100 | (6, 7, 9, 10, 91) |
| Total SPEN abundance | *S*<sub>*T*</sub> | [10<sup>1.5</sup>,10<sup>4.5</sup>] | 1000 molecules | 70 molecules | 250 | (87–90) |
| Maximum number of SPEN that can bind to an *Xist* molecule | *N*<sub>*S*</sub> | $round(\frac{S_T}{X_{bs}})$ | 10 molecules | 1 molecule | 3 molecules | (7) |

Each X chromosome contains a gene that codes for *Xist*. We assume *Xist* expression is induced by a transcriptional activator that is expressed from each X chromosome and acts in trans, creating a common pool of the activator in the nucleus. We assume that because of *Xist*’s slow rate of diffusion compared to its half-life, *Xist* acts predominantly *in cis*, associating almost exclusively with the X chromosome from which it was expressed, an assumption that is generally supported by experimental observations (7, 10, 14, 44, 50, 51). However, in the supplement we relax this assumption and show that it is not necessary for *Xist* to act exclusively in cis for our model to function properly. Once expressed, *Xist* recruits the RNA-binding protein SPEN that inhibits expression of all X-linked genes including the activator for *Xist* (11–13, 17, 18, 22, 23). Moreover, because SPEN has been shown to promote *Xist* stability in embryonic stem cells (10, 52), we assumed that SPEN stabilizes *Xist*’s association with the X chromosome and slows its rate of degradation. Previous studies showed that SPEN aggregates in supramolecular complexes (SMACs) nucleated by *Xist* through interaction of SPEN’s intrinsically disordered region (IDR) (7, 22). Each SMAC consists of around 35 copies of SPEN (7). Therefore, we assumed one *Xist* molecule can recruit multiple SPEN proteins. Thus, in our model, interactions between *Xist* and SPEN form a positive feedback loop which we hypothesize produces an *Xist*-SPEN assembly sufficient to cover the X chromosome. As SPEN accumulates around the X chromosome, it eventually silences expression of the activator forming a negative feedback loop. Below, we use the term “bound” to refer to stable association of *Xist* or SPEN with X-linked chromatin, and the term “free” to refer to diffusible *Xist* or SPEN.

To investigate the behavior of the model we cast it as a set of coupled ordinary differential equations (ODEs) for the following variables: 1) X_bi_ – number of *Xist* bound to the ith X chromosome, 2) X_fi_ – abundance of free *Xist* associated with the ith X chromosome, 3) S_bi_ –number of SPEN bound to the ith X chromosome, and 4) A_x_ – abundance of the trans-acting activator for *Xist*. We assume that the total amount of SPEN is conserved and is given by

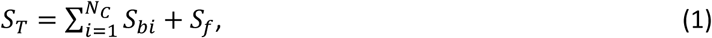

where N_c_ is the total number of X chromosomes and S_f_ is the abundance of free SPEN.

The equation for X_fi_ is given by

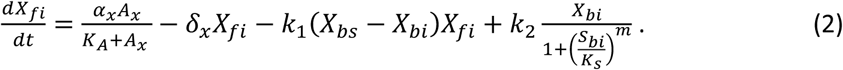

The first term on the right-hand-side of Eq. (2) represents the induction of *Xist* by A_x_. In this term α_x_ is the maximum synthesis rate of *Xist* and *K_A_* is the level of the activator for which the synthesis rate is half its max. The second term represents the degradation of free *Xist*. The third term represents the binding of free *Xist* to the i^th^ chromosome. In this term X_bs_ represents the total number of *Xist* binding sites on an X chromosome and k_1_ is the rate constant for association. The last term represents the dissociation of *Xist* from the i^th^ X chromosome. The parameter k_2_ is the rate constant for the dissociation of *Xist* in the absence of bound SPEN. The denominator in this term models the effect of SPEN stabilizing *Xist*’s association with the X chromosome.

The equation for the number of *Xist* bound to the i^th^ X chromosome is

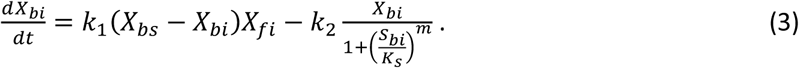

The equation for the number of SPEN bound to the i^th^ X chromosome is

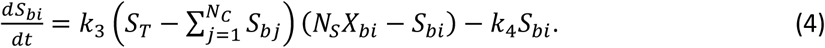

The first term on the right-hand-side of Eq. (4) represents SPEN binding to *Xist* molecules associated with the i^th^ X chromosome. The parameter k_3_ is the rate constant for SPEN binding. The quantity in the first set of parentheses is the amount of free SPEN, and the term in the second set represents the number of free SPEN binding sites associated with bound *Xist* molecules with N_S_ being the number of binding sites per *Xist* molecule. The last term in the above equation represents the dissociation of bound SPEN, where k_4_ is the dissociation rate constant.

The equation for the abundance of the activator A_x_ is given by

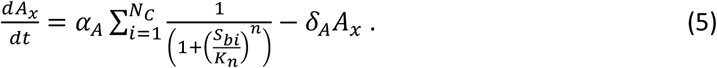

The first term on the right-hand side of Eq. (5) represents the synthesis rate for A_x_. In this term α_A_ is the constitutive synthesis rate. The Hill functions in the summation capture repression by SPEN bound to the j^th^ X chromosome. The last term in the equation represents degradation of A_x_. A description of the model parameters and the values used in the results are given in Table 1.

### Two non-exclusive mechanisms produce the n-1 rule

To determine if the model given by Eqs. (1–5) can produce results consistent with experimental observations, we used an evolutionary algorithm (EA) to search parameter space (53, 54). To implement the EA, we constructed an objective function that is minimized by parameter sets that reproduce three critical properties of X chromosome inactivation (see Supporting Information for details): 1) The X chromosome in males remains active. 2) Females inactivate one X chromosome. 3) Female cells with an extra copy of the X chromosome inactivate two X chromosomes. For parameter sets to be considered successful they must produce model behavior that satisfies all three properties. Bounds included in the EA for the model parameters were carefully selected to be consistent with previous experimental studies of XCI (Supporting Information).

We initially included a constraint in the objective function that penalized solutions with high levels of bound SPEN. The rationale for including this term came from our preliminary investigations in which we tuned parameters by hand. These simulations suggested that a mechanism that relies on SPEN depletion would have difficulty generating the n-1 rule for cells that contained greater than two X chromosomes because competition for free SPEN typically produced a “winner-take-all” scenario with a single X chromosome being silenced. This constraint can also be rationalized biologically, in the sense that there is likely to be a limit on the amount of SPEN that can be complexed with a single molecule of *Xist*. As described below, we also ran the EA without this constraint to search for additional mechanisms that might underlie XCI.

We ran the EA for 501 realizations on populations of size 500 with 100 generations in each realization. This resulted in 36 parameter sets satisfying all three criteria (Fig. 1B). We ran further simulations of the successful parameter sets to confirm that the results held regardless of the initial conditions (Fig. S1). In all successful parameter sets, the Hill coefficient *m* in the term that models SPEN’s stabilization of *Xist*’s interaction with the X chromosome is greater than one. This suggests that positive feedback between *Xist* and SPEN is critical for XCI. The values for the Hill constant for this term, K_s_, all lie between 1 and 10, indicating that in our model, binding of only a few SPEN molecules is sufficient to stabilize *Xist* on the X chromosome.

Typical results for male and female cells (XX and XXX) are shown in Figs. 1C-E. In XX female cells, the model can be explained as follows. Before X chromosome inactivation occurs, the activator is expressed from both X chromosomes, and its abundance reaches a level that promotes transcription of *Xist* from both X chromosomes. However, owing to natural stochasticity in the transcriptional process, the two X chromosomes will not produce exactly same number of molecules of *Xist* at any given time. Once produced, *Xist* binds to the X chromosomes and recruits SPEN. Positive feedback between *Xist* and SPEN (i.e., *Xist* recruiting SPEN and SPEN stabilizing *Xist*) causes a rapid accumulation of these molecules on both X chromosomes (Fig. 1D). Accumulation of SPEN inhibits activator production that in turn decreases the synthesis rate of *Xist* causing the levels of bound SPEN and *Xist* to drop on both X chromosomes. The positive feedback loop is broken on the X chromosome with fewest *Xist* molecules, causing a rapid loss of bound *Xist* and SPEN from this chromosome (Fig. 1D, blue curves). The decrease in bound SPEN restores activator expression from this X chromosome that in turn increases the *Xist* transcription rate on both X chromosomes. However, with activator expression from only a single X chromosome, the *Xist*/SPEN positive feedback loop cannot be reinitiated on the X chromosome with fewest bound *Xist* but is sustained on the other X chromosome (Fig. 1D, red curves). This illustrates an important feature of the model that higher levels of activator are needed to initiate the *Xist*/SPEN positive feedback loop than sustain it, a general feature of positive feedback that has been observed previously in many systems (55). Similarly, in male cells (XY), activator expression is again insufficient to exceed the threshold necessary to initiate XIST/SPEN positive feedback (Fig. 1C). Furthermore, if in male cells the threshold were transiently exceeded, positive feedback could not be sustained because as SPEN accumulated on the single X chromosome, expression of the activator would drop below the level required to sustain XCI. Note that at steady state, there are roughly equal levels of activator expressed in male and female cells, because in both cases activator expression is driven from a single X-chromosome. For the case when 3 X chromosomes are present, the model works similarly to the 2X case (Fig. 1E).

We wondered if other mechanisms could capture the three criteria for XCI. To test this, we removed the constraint in the objective function that penalized solutions with high levels of bound SPEN. Surprisingly, the EA found parameter sets that captured all three criteria and relied on SPEN depletion to satisfy the n-1 rule. No other mechanisms for XCI were found by the EA. The fact that the EA did not generate other mechanisms suggests that if they do exist, they occupy a relatively small region of parameter space, making them less likely to occur.

To further investigate the SPEN depletion mechanism, we updated the objective function to penalize solutions with high levels of free SPEN following XCI (see Methods). In the SPEN depletion mechanism, male cells (XY) still do not inactivate their single X chromosome because similar to the activator inhibitor model activator expression from a single X chromosome is insufficient to initiate the *Xist*/SPEN positive feedback loop (Fig. 2A), and for female cells (XX) positive feedback between *Xist* and SPEN results in a winner-take-all scenario: the X chromosome with higher levels of bound *Xist* and SPEN outcompetes the other and depletes the pool of free SPEN (Fig. 2B). The pool of free SPEN reaches its steady-state concentration when its association rate for binding to either chromosome is balanced by the rate of dissociation from either chromosome. Because of the high affinity of SPEN binding to Xist in this model, at steady state the fraction of free SPEN is small. Depletion of free SPEN relieves repression of the activator on the other X chromosome, causing activator levels to rise.

**Figure 2.**
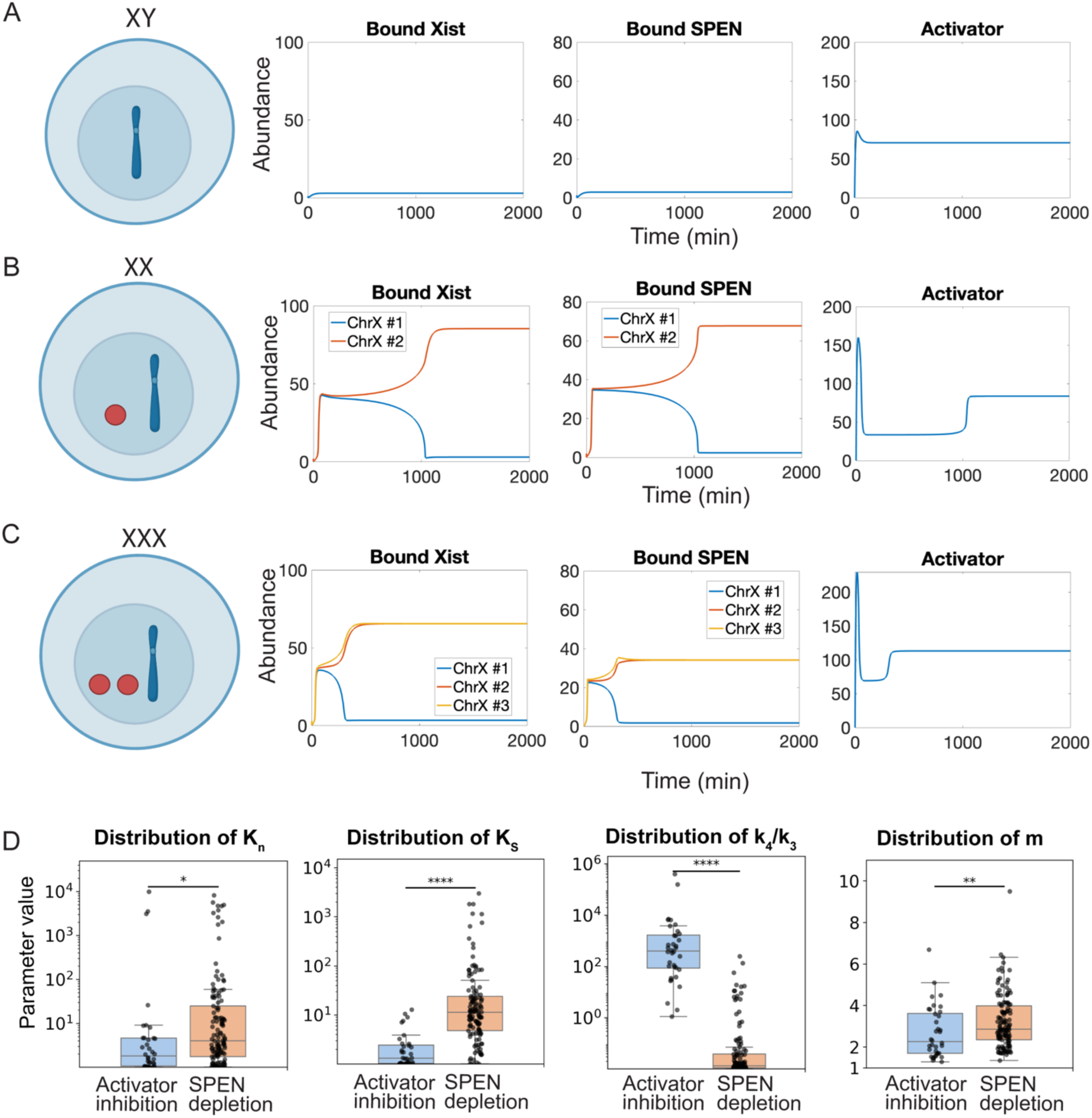
SPEN depletion model for XCI. **A**) Time series for number of molecules of bound *Xist*, bound SPEN, and activator (i.e. Abundance) for XY males. The total amount of SPEN molecules per cell in each simulation is S_T_ = 70. **B**) Same as **A** except for the case of XX females. **C**) Same as **A** except for XXX females. **D**) Distributions of parameters that are significantly different between the models: K_n_ = # of SPEN molecules for which activator production rate is ½ max, K_s_ = # of SPEN molecules at which *Xist* dissociation rate is ½ max, k_4_/k_3_ ratio of SPEN dissociation rate to association rate, and m Hill coefficient for SPEN inhibition of *Xist* dissociation (* p ≤ 0.05, ** p ≤ 0.01, *** p ≤ 0.001, **** p ≤ 0.0001). Y axes of the first three panels are log_10_. Y axis of the last panel is linear.

Surprisingly, this mechanism also adheres to the n-1 rule for cases where n > 2, instead of resulting in a single X chromosome being silenced (Fig. 2C). When more than one X chromosome is active, the activator level is high enough to drive *Xist* expression on multiple chromosomes that in turn, compete for SPEN. Because only a finite number of SPEN can be associated with an X chromosome (limited by the number of *Xist* binding sites on the chromosome and number of SPEN that can bind to an *Xist* molecule), the ability of X chromosomes to recruit SPEN is reduced by SPEN accumulation. This allows X chromosomes with fewer bound SPEN to out compete those with more bound SPEN for free SPEN, causing bound SPEN levels to equalize on inactivated X chromosomes. Recruitment of SPEN continues until virtually all the SPEN is evenly distributed between n-1 chromosomes. Expression of the activator from the remaining active X chromosome is derepressed and the activator level rises (Fig. 2C). However, this remaining X chromosome cannot compete with the n-1 silenced chromosomes for the small remaining pool of free SPEN, and therefore, remains active. We conducted additional simulations with varying initial levels of bound *Xist* to confirm the n-1 rule holds regardless of initial conditions (Fig. S2).

The parameter distributions for the two mechanisms have the following differences. For the activator inhibition mechanism, a smaller amount of bound SPEN is required to repress production of the activator and stabilize *Xist* on the X chromosome as indicated by the smaller values of K_n_ and K_S_ for the activator inhibition model (Fig. 2D, left two panels). SPEN and *Xist* have a higher affinity (smaller k_4_/k_3_) in the SPEN depletion mechanism than the activator inhibition mechanism (Fig. 2D, third panel) and larger Hill coefficient for the effect of SPEN on the dissociation rate of *Xist* (Fig. 2D, right panel). These differences allow the concentration of free SPEN to be efficiently depleted in the SPEN depletion mechanism.

The form of positive feedback loop between SPEN and *Xist* is not limited to the stabilization of *Xist* by SPEN. We also investigated a feedback mechanism in which SPEN actively promotes the binding of *Xist* to the chromosome, which creates a mutual activation loop. In this loop, SPEN enhances the recruitment of *Xist* on the chromosome and further amplifies accumulation of SPEN. This mechanism produced similar results for both models (Figs. S3 and S4). These findings demonstrate that in our models, the precise molecular details of the feedback loop can vary, but the core principle of positive feedback between *Xist* and SPEN is required to drive the symmetry breaking for XCI.

### XCI in polyploidy cells

In tetraploid (XXXX) cells, it has been observed that 2 of the 4 X chromosomes are inactive (4). Tetraploid cells also tend to have larger cytoplasmic and nuclear volumes (56). One potential effect of increased nuclear volume is a dilution in the concentrations of *Xist*, SPEN and the activator. To test if dilution effects were sufficient to switch from 3 to 2 silenced X chromosomes, we focused on the activator inhibition model. Eqs. (1–5) are written in terms of molecular abundances for which the synthesis rates, α_x_ and α_A_, have units of per unit time. If we assume that these rates remain constant as the volume is increased, then we expect the concentrations of *Xist* and the activator to be reduced. Similarly, we assume that the molecular abundance of SPEN remains constant, and therefore the concentration is reduced with increased volume. Because we are using molecular abundances as opposed to concentrations in our model, we scale second order rate constants as 1/volume to account for a change in volume. To initially investigate the behavior of the activator inhibition model as the nuclear volume changes with four X chromosomes, we created a 2-parameter bifurcation diagram for the second order rate constants (Fig. 3A). We found parameter regimes with 0 – 3 inactive X chromosomes. We used these results to select a parameter set for which the number of silenced chromosomes decreased as the volume increased (Fig. 3B). Our model predicts that roughly a 15-40 percent increase in the nuclear volume is sufficient to explain the behavior of tetraploid female cells, and that increases in volume beyond this range would cause failure of the n-1 rule (Fig. 3B).

**Figure 3.**
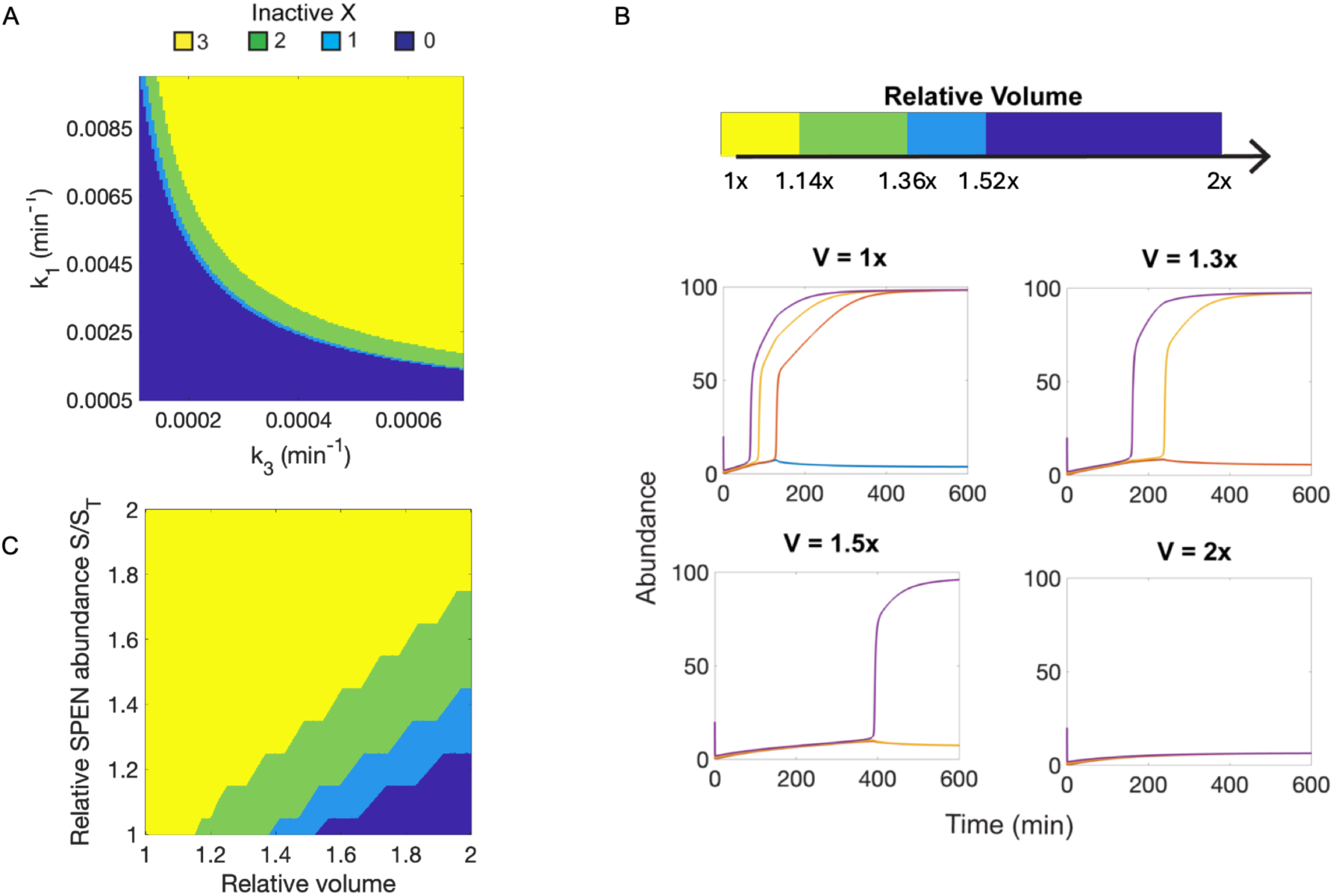
Results for tetraploid (XXXX) females using the activator inhibition model. **A**) Two-parameter bifurcation diagram using the *Xist*-chromosome binding rate *k*_1_ and the SPEN-*Xist* binding rate *k*_3_as bifurcation parameters. Other parameter values are given in Table 1. The regime in which 2 of the 4 X-chromosomes are silenced is marked in green. **B)** Number of silenced X chromosomes as a function of the relative nuclear volume. Each X chromosome is represented by a different colored line. Representative time series for bound Xist molecules are also presented for each regime. In each simulation, the initial number of molecules of *Xist* bound to each of the four X-chromosomes was 1, 2, 10, and 20, respectively. **C)** Number of inactive X chromosomes when varying relative nuclear volume and SPEN abundance. Original abundance of SPEN (S_T_) is equal to 1000.

Recent experiments in yeast revealed that protein abundances both increase and decrease with ploidy with the average trend being a sublinear increase (57). To investigate how the model would function if levels of total SPEN were allowed to increase, we ran simulations in which the total SPEN abundance and nuclear volume were varied (Fig. 3C). These simulations revealed that for a 35% increase nuclear volume the model predicts a 21% increase total SPEN can be tolerated before 3 instead of 2 X chromosomes are silenced. If the nuclear volume doubled, the total SPEN must increase between 45-74% to insure two X chromosomes are silenced in tetraploid cells.

### Transient expression of *Xist* in males

*Xist* is transiently expressed on the single X chromosome in males (33–38, 58, 59). A search of our parameter sets revealed such behavior for both models, although it occurred rarely. Therefore, to select for this behavior and enable its study, we added a 4^th^ term to our objective function that selected for solutions in which the maximum amount of bound *Xist* is greater than its steady state value for the case of male cells (Methods). We then reran the EA for the activator inhibitor model. The EA generated parameter sets that produced transient accumulation of *Xist* in male cells and captured the other properties of XCI (Fig 4). However, an analysis of the model revealed that it produced multiple, stable steady states, depending on the initial conditions. For the case of XX cells, steady states with either a single X chromosome silenced or no X chromosome silenced were observed, and for the case of XXX cells, steady states with 0, 1 or 2 X chromosomes silenced were observed.

**Figure 4.**
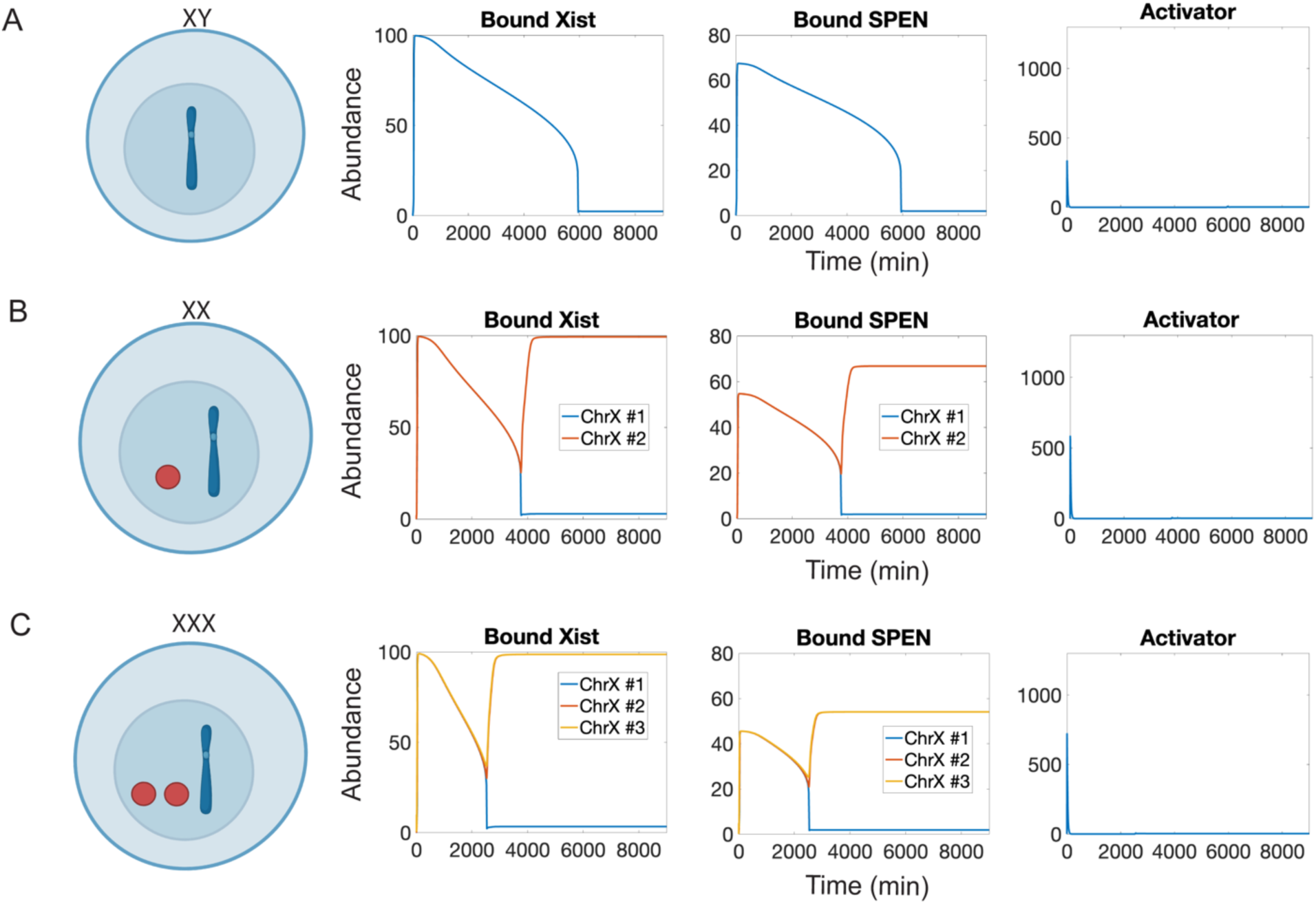
Transient *Xist* expression in males for the activator inhibition model. **A**) Time series for the number of molecules of bound *Xist*, bound SPEN, and activator (i.e. Abundance), showing transient *Xist* expression in males. The total amount of SPEN molecules per cell in each simulation is S_T_ = 250. **B**) Same as **A** except for the case XX females. **C**) Same as **A** except for XXX females.

To quantify the frequency at which these differing outcomes occurred in XX and XXX cells, we ran 10,000 simulations for each case using random initial conditions drawn from uniform distributions that spanned the relevant ranges for the model variables. For XX cells less than 1% of the simulations did not lead to a proper n-1 XCI and for the case of XXX less than 4% of the simulations did not evolve to a solution with the expected number of two X chromosomes silenced. Given the small percentage of times the model produced the incorrect behavior, these states could be classified as “unlikely”, and if they did occur in vivo, would presumably lead to a tolerable level of selection against cells with failed XCI. It is also possible other regulatory mechanisms not included in the model could further insure XCI is correctly achieved. Considering the parameter values that generate transient *Xist* expression in males versus the ones used in Fig. 1 that do not, the largest differences in the former are that the synthesis rate of the activator is faster (66.8 min^-1^ versus 1.1 min^-1^), the degradation rate of the activator is slower (0.048 min^-1^ versus 0.72 min^-1^) and the association rate for SPEN binding to *Xist* is faster (0.014 min^-1^ versus 2.6 x 10^-4^ min^-1^). These differences allow the *Xist*/SPEN positive feedback loop initially to be activated in males and transiently maintain bound molecules of *Xist* after the activator level falls.

### Particle-based model for X-chromosome inactivation

We chose initially to formulate our models in terms of ODEs because of the relative mathematical simplicity of ODEs and the efficiency with which they can be simulated. However, ODE models fail to capture spatial variations in concentrations or the stochastic nature of biochemical reactions. To capture these effects, we developed a particle-based model in which individual molecules of *Xist,* SPEN, and the activator are tracked as they diffuse and react in the nucleus. X chromosomes were assumed to occupy spherical domains of radius 2.5 µm. Within these domains, 100 *Xist* binding sites were randomly placed using a 3D random walk (Fig. 5A) (Supporting Information). Activator proteins bind to the X chromosome at the site where *Xist* is synthesized, and once bound, increase the synthesis rate of *Xist*. SPEN was modeled as binding to *Xist* only when the latter was bound to X chromosomes. Repression of the activator by SPEN was modeled using the synthesis rate in Eq. (5). We sought to construct our model to be consistent with data that suggest that SPEN and *Xist* aggregate in supramolecular complexes along the X chromosome and stabilize each other, either through direct contacts or multivalent SPEN-SPEN interactions (7, 10, 22). Simulating the multivalent interactions involved in forming condensates is computationally intensive (60, 61). Therefore, similar to the ODE model, we did not explicitly consider SPEN-SPEN interactions but rather assumed that the dissociation rate of *Xist* from an X chromosome depends on the overall number of SPEN molecules bound to the chromosome. We used the parameter values from the ODE model to estimate rates in the particle-based simulations and then tuned the parameters by hand. Parameters values used in the particle-based simulations are given in Table 2 and full details for the simulations are presented in the Supporting Information.

**Figure 5.**
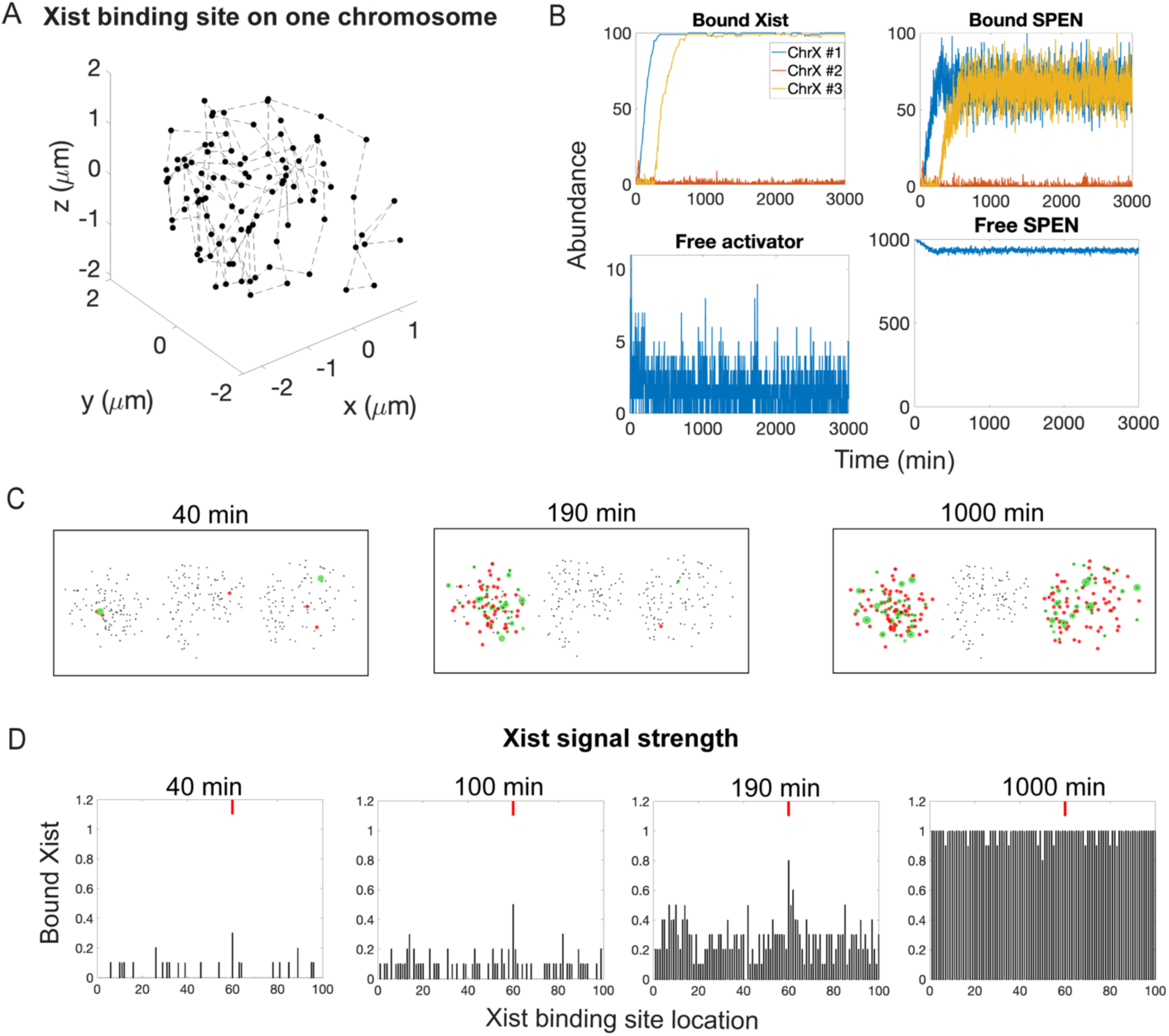
Particle-based model for XCI. **A)** Example for *Xist* binding sites generated by a 3D random walk placed with step size 1 *μ*m. **B)** Time series for the molecular abundance of each species in the model for the case of XXX females. **C)** Snap shots showing simulation results at different times (black dots – *Xist* binding sites, red dots – bound molecules of *Xist* and green dots *Xist*-SPEN complex). **D**) Fraction of each of 100 sites along an inactive X-chromosome that are bound by *Xist* across 10 simulations, at time points of 40, 100, 190, and 1000 min. The red lines denote the *Xist* transcription site.

**Table 2.** Summary of Particle-based Model Parameters.

| Parameter description | Symbol | Parameter values for Particle-based model |
|---|---|---|
| *Xist*-chromosome association rate | *k*<sub>1</sub> | 250 min<sup>-1</sup> |
| *Xist*-chromosome dissociation rate in the absence of SPEN | *k*<sub>2</sub> | 8.7 min<sup>-1</sup> |
| SPEN-*Xist* association rate | *k*<sub>3</sub> | 5000 min<sup>-1</sup> |
| SPEN-*Xist* dissociation rate | *k*<sub>4</sub> | 10.2 min<sup>-1</sup> |
| Maximum *Xist* transcription rate | *α*<sub>*x*</sub> | 3.2 min<sup>-1</sup> |
| Degradation rate for free *Xist* | *δ*<sub>*x*</sub> | 0.0076 min<sup>-1</sup> |
| Maximum activator production rate | *α*<sub>*A*</sub> | 1.1 min<sup>-1</sup> |
| Degradation rate for the activator | *δ*<sub>*A*</sub> | 0.72 min<sup>-1</sup> |
| Number of activator molecules at which the *Xist* transcription rate is half max | *K*<sub>*A*</sub> | 16.1 molecules |
| Number of SPEN molecules at which the *Xist* dissociation rate is half max | *K*<sub>*S*</sub> | 2.06 molecules |
| Hill coefficient for SPEN inhibition of *Xist* dissociation | m | 3.09 |
| Number of SPEN molecules at which the activator production rate is half max | *K*<sub>*n*</sub> | 2.2 molecules |
| Hill coefficient for SPEN inhibition of activator production | n | 9.4 |
| Total *Xist* binding sites per chromosome | *X*<sub>*bs*</sub> | 100 molecules |
| Total SPEN abundance | *S*<sub>*T*</sub> | 1000 molecules |
| *Xist* diffusion coefficient | *D*<sub>*x*</sub> | 0.4 *μ*m<sup>2</sup> min<sup>-1</sup> |
| Activator diffusion coefficient | *D*<sub>*A*</sub> | 15 *μ*m<sup>2</sup> min<sup>-1</sup> |
| SPEN diffusion coefficient | *D*<sub>*S*</sub> | 15 *μ*m<sup>2</sup> min<sup>-1</sup> |
| Maximum number of SPEN that can bind to an *Xist* molecule | *N*<sub>*S*</sub> | 10 molecules |
| Reaction radius | *ρ* | 0.1 μm |
| Time step | *δt* | 0.0002 min |

We used our particle-based model to simulate normal female (XX) and male (XY) (Fig. S5) and female (XXX) cells and found that the results were strikingly concordant with expectations based on experimental evidence (Figs. 5B-C, Movies S1-S3). For each genotype, the particle-based simulation reproduced the n-1 rule, and for genotypes with more than 1 X chromosome, the silenced chromosomes were randomly selected in a manner determined by intrinsic fluctuation, consistent with observations in cells (4). The time scale for XCI to occur was on the order of 16 hrs, a number within range of in vivo estimates (18, 32, 62). Moreover, *Xist* initially accumulated closest to the site of transcription then spread until the X chromosome was uniformly covered, an observation also consistent with in vivo measurements (Fig. 5D; (63, 64)). Lastly, in all simulations, once an X chromosome was silenced it remained silenced for the duration of the simulation, consistent with the fundamental outcome of XCI.

## Discussion

Three important features of XCI are: 1) Symmetry breaking: X chromosomes are selected at random for inactivation, 2) Counting rules: n–1 X chromosomes in diploid cells and n–2 in tetraploid cells are inactivated, and 3) Robustness: XCI is reliable despite stochasticity in biochemical processes and variability across individuals. We demonstrate that mathematical models that rely on the integration of positive and negative feedback loops can capture each of these features. In our models, a positive feedback loop -- formed by SPEN’s stabilization of *Xist* on X chromosomes and *Xist*’s ability to recruit multiple SPEN molecules -- provides a mechanism to amplify small differences between X chromosomes, causes *Xist*-SPEN complexes to cover the X chromosomes in an all-or-none fashion, and thereby establishes symmetry breaking. Conversely, a negative feedback loop -- formed by SPEN’s repression of an *Xist* activator -- enforces the n-1 rule by limiting the number of chromosomes that express sufficient *Xist* to exceed the threshold needed to initiate *Xist*/SPEN positive feedback.

Applying these positive and negative feedback loops in different parameter regimes, we found two mechanisms that can successfully recapitulate most if not all known chromosomal counting scenarios that result in XCI. In the first mechanism, SPEN silences all but one of the X chromosomes, at which point the activator concentration drops below the level required to trigger XCI. In the second mechanism, positive feedback produces competition for SPEN between X chromosomes and depletion of SPEN ensures all but one X chromosome is inactivated. These models offer new perspectives on symmetry-breaking and XCI that await experimental interrogation.

Our study focused on SPEN as the sole entity required for *Xist*-induced silencing (10–19). However, the concept of SPEN in our models could be viewed a proxy for the minimal set of proteins required for XCI. Multiple factors, including the Polycomb Repressive Complexes, are also required, and evidence suggests that even SPEN itself requires cofactors for recruitment to *Xis*t (24–32). Thus, in its complete form, the silencing-competent *Xis*t ribonucleoprotein complex includes components in addition to SPEN, each of which carries on- and off-rates and potential to engage in feedback.

### Comparison with existing models

A mathematical model describing the onset of random XCI has been proposed (35). Like our model above, this model also includes a threshold concentration for a trans-acting *Xist* activator that cannot be met by a single X chromosome. However, to break and subsequently maintain asymmetric expression of *Xist*, it additionally relies on mutual inhibition between *Xist* and an X-linked repressor of *Xist*, which in the mouse, was proposed to be the antisense lncRNA *Tsix*. The model notably recapitulates essentially all experimental observations and clearly represents a viable solution to the n-1 rule of XCI (35). Still, it is worth noting that the mechanisms responsible for the monoallelic expression of *Xist* remain incompletely defined, and while many experimental studies of XCI have been performed in mice, *Xist* is embedded in a complex cis-regulatory landscape that varies dramatically between eutherians. For example, the murine lncRNA *Tsix* does not exhibit strong evidence of conservation in humans and its role in the establishment of *Xist* monoallelism in the mouse has been questioned (41, 65, 66). Thus, it seems reasonable that additional models to explain the n-1 rule of XCI may exist.

Our model is distinct in that it does not require a cis-acting repressor of *Xist* or even competition between different cis-acting DNA regulatory elements to achieve monoallelic expression of *Xist*. Rather, *Xist* monoallelism is achieved using a positive feedback loop that results in stabilization of *Xist* by a common pool of SPEN. We envision that symmetry between X chromosomes could be broken through random fluctuations in *Xist* expression, which would occur as the result of bursts of activity that are inherent to the transcriptional process (67–69). In our model, whichever X chromosome experiences larger bursts of *Xist* transcription during the early stages of XCI is the chromosome that will eventually outcompete the other when levels of the X-linked *Xist* activator or SPEN become limiting. This model obviates the need for competition between cis-acting DNA elements and trans-acting factors and instead underscores how the asymmetric propagation of an RNA-protein effector complex might play an important role in establishing the monoallelic expression of *Xist*.

A body of work indicates that *Xist* is embedded in a complex cis-regulatory landscape that includes its own promoter, DNA enhancer elements, and multiple lncRNAs organized into discrete topological domains, and that together, these entities play important roles in producing and enforcing *Xist*’s monoallelic expression (39–41). Still, the success of our model, its independence from multi-element cis-regulatory circuits, and its relative simplicity leads us to speculate that symmetry breaking during ancestral forms of *Xist*-dependent XCI in eutherians may have been driven by an inherent ability of the *Xist* ribonucleoprotein complex to propagate asymmetrically. Under this model, the multicomponent DNA regulatory landscape that surrounds the *Xist* locus may have evolved after the emergence of XCI rather than as a precursor to it, to optimize the magnitude, duration, and frequency of bursts of *Xist* transcription. Beyond putative effects on bursting, it is possible that another major function of *Xist*’s cis-regulatory landscape may be to maintain asymmetry between the X chromosomes after XCI has proceeded to a semi-stable state, ensuring that *Xist* expression is kept off on the active X chromosome in differentiated progeny. The discovery that *Tsix*, a repressor of *Xist* expression in mice, functions to maintain *Xist* monoallelism rather than establish it, is consistent with this latter idea (65, 66).

In considering *Xist* as an essential component of its own monoallelic expression system, our study provides new perspective to additional experimental observations. In most eutherians, including humans and primates, both X chromosomes in females frequently accumulate *Xist* “clouds” and are enriched in silencing-associated chromatin modifications (presumably over non-transcribed regions) prior to the induction of gene silencing and bona fide XCI (4, 33–38). Thus, it is only upon the activation of its gene silencing function that *Xist* transitions to a monoallelic state. This transition has been observed to occur stochastically in cells (4), and would be predicted by our model, under the assumption that at some point during the silencing process, the two X chromosomes will harbor stochastic differences in the numbers of bound *Xist* molecules.

### Broader i mplications and future directions

Our model has broader implications for understanding symmetry breaking in biological systems. Examples of symmetry breaking include establishment of the body’s left-right asymmetry, asymmetric cell division, axon selection in neurons, and cellular polarity establishment. A common mechanism for symmetry breaking is positive feedback to amplify slight differences in chemical concentrations that occur due to intrinsic fluctuations. A key difference between symmetry breaking during XCI and other symmetry breaking phenomena, such as axon selection or polarity establishment, is that the latter two processes follow a “winner-take-all” scenario. That is, initially multiple neurites or potential polarity sites exist. However, positive feedback ensures that once a neurite or nascent polarity site gains an advantage, it out competes the others for a shared pool of molecular components. This competition ensures a single axon or polarity site is formed. In contrast, during XCI there is a single “loser”. That is, only 1 X chromosome is unable to recruit SPEN and remains active. Our model implements this single loser scenario by ensuring that a single X chromosome is insufficient to activate the *Xist*/SPEN positive feedback loop. The inactivation of all but one X chromosome versus the creation of a unique active site requires an additional level of regulation. For example, in polarity establishment, a single positive feedback loop is sufficient to ensure the formation of a single polarity site (70). However, XCI requires a positive and negative feedback loop. Positive feedback generates multiple steady states (e.g., female cells have 2 steady states represented by 1 of the 2 X chromosomes becoming silenced), and negative feedback ensures the n-1 rule applies for cases with n > 2.

Evidence suggests that RNA-protein effector complexes promote monoallelism at olfactory receptor loci (71). Here, the data can be described by a “winner-take-all” mechanism where RNA molecules enable a single regulatory hub to outcompete others for a limited pool of trans-activator. This contrasts with our model where there is a single “loser” chromosome that cannot recruit enough SPEN to become silenced. We envision that single-gene, monoallelism on autosomes may be more frequently characterized by winner-take-all events, which mechanistically, require fewer steps to evolve. Genes controlled by transcriptional activators that are prone to accumulating into ribonucleoprotein assemblies might frequently experience monoallelism, under a model where RNA-driven assemblies distribute the activators in a way that makes their target genes prone to all-or-none expression. RNA elements within a gene (or nearby enhancer or lncRNA) that promote partitioning of transcriptional activators into RNA-seeded assemblies might increase the gene’s frequency of monoallelism to the extent that it occurs reproducibly (40, 72–74). Therefore, it would be interesting to consider possible roles for RNA-protein assemblies in driving monoallelic expression events on autosomes, which are widespread (40, 72–74).

A difference between our model and the one proposed in (35) is that we predict that allelic differences in the amount of SPEN bound to *Xist* determine which chromosome ultimately becomes inactivated. Under this prediction, monoallelic deletion of the SPEN-binding region of *Xist* would bias XCI towards the X chromosome expressing wild-type *Xist*, and homozygous deletion of SPEN would prevent monoallelic *Xist* from occurring. These experiments have been performed in mouse embryonic stem cell models, showing results at least partially consistent with our predictions, but with interpretations complicated by the presence of *Tsix*, which plays a role in *Xist* repression predominantly after differentiation of the epiblast (52, 66, 75). Therefore, investigating the role of SPEN in symmetry breaking in human model systems, which appear to lack a functional *Tsix*, or in mouse models prior to differentiation of the epiblast, before *Tsix* exerts its repressive effect (52, 66, 76–79) will provide important tests moving forward.

Another potential avenue for testing predictions of our model is the cis-acting polymorphic locus termed the X controlling element (*Xce*), which affects the frequency with which individual X chromosomes are selected for inactivation. In hybrids, stronger *Xce* alleles paired with weaker ones result in preferential inactivation of the X chromosome harboring the weaker allele (80). Because in our model the choice of which X chromosome is inactivated is driven by stochastic differences in the numbers of bound *Xist* molecules, we predict that *Xce* alleles exert their effects by modulating the frequency, duration, or magnitude of *Xist* transcriptional bursts during early stages of XCI. These features could be quantitatively measured for different *Xce* alleles to determine if they correlate with the strength of the allele for biasing XCI.

## Materials and Methods

### ODE Simulations

The ODE models were simulated using MATLAB’s (R2024a) built-in function *ode15s* and validated using *ode45*. To optimize the parameters in our model, we adapted the evolutionary algorithm (EA) from Pomeroy et al. that uses DEAP (Distributed Evolutionary Algorithms in Python) (54, 81, 82). Full details can be found in the Supporting Information.

### Particle-based Simulations

The particle-based model was simulated using Smoldyn (v3.71) (83), on a Linux-based computing system (Longleaf cluster at UNC Chapel Hill). Smoldyn treats all molecules as point particles undergoing Brownian motion, with their stochastic trajectories integrated using the Euler–Maruyama method. Details for the particle-based simulations are presented in the Supporting Information.

All code is available at https://github.com/guanhighyun/XCI_model

## Acknowledgments

We thank Sundeep Kalantry for his critical reading of the manuscript and helpful suggestions to improve clarity. Portions of the paper were developed from the thesis of Kaiyun Guan. Model schematics and chromosome diagrams were created with BioRender.com. This Study was supported by NIH Grants R35GM127145 (TCE) and R35GM153293 (JMC).

## References

1. F. Dossin, E. Heard, The Molecular and Nuclear Dynamics of X-Chromosome Inactivation. Cold Spring Harb Perspect Biol 14, a040196 (2022).

2. C. J. Brown, et al., A gene from the region of the human X inactivation centre is expressed exclusively from the inactive X chromosome. Nature 349, 38–44 (1991).

3. M. Lyon, X-chromosome inactivation and human genetic disease. Acta Paediatrica 91, 107–112 (2002).

4. K. Monkhorst, I. Jonkers, E. Rentmeester, F. Grosveld, J. Gribnau, X Inactivation Counting and Choice Is a Stochastic Process: Evidence for Involvement of an X-Linked Activator. Cell 132, 410–421 (2008).

5. S. Webb, T. J. de Vries, M. H. Kaufman, The differential staining pattern of the X chromosome in the embryonic and extraembryonic tissues of postimplantation homozygous tetraploid mouse embryos. Genet Res 59, 205–214 (1992).

6. A. Cerase, et al., Spatial separation of Xist RNA and polycomb proteins revealed by superresolution microscopy. Proceedings of the National Academy of Sciences 111, 2235–2240 (2014).

7. Y. Markaki, et al., Xist nucleates local protein gradients to propagate silencing across the X chromosome. Cell 184, 6212 (2021).

8. M. D. Schertzer, et al., lncRNA-Induced Spread of Polycomb Controlled by Genome Architecture, RNA Abundance, and CpG Island DNA. Mol Cell 75, 523–537.e10 (2019).

9. H. Sunwoo, J. Y. Wu, J. T. Lee, The Xist RNA-PRC2 complex at 20-nm resolution reveals a low Xist stoichiometry and suggests a hit-and-run mechanism in mouse cells. Proceedings of the National Academy of Sciences 112, E4216–E4225 (2015).

10. L. Rodermund, et al., Time-resolved structured illumination microscopy reveals key principles of Xist RNA spreading. Science 372, eabe7500 (2021).

11. C. A. McHugh, et al., The Xist lncRNA directly interacts with SHARP to silence transcription through HDAC3. Nature 521, 232–236 (2015).

12. C. Chu, et al., Systematic discovery of Xist RNA binding proteins. Cell 161, 404–416 (2015).

13. B. Moindrot, et al., A Pooled shRNA Screen Identifies Rbm15, Spen, and Wtap as Factors Required for Xist RNA-Mediated Silencing. Cell Rep 12, 562–572 (2015).

14. I. Dror, et al., XIST directly regulates X-linked and autosomal genes in naive human pluripotent cells. Cell 187, 110–129.e31 (2024).

15. C. Alfeghaly, et al., XIST dampens X chromosome activity in a SPEN-dependent manner during early human development. Nat Struct Mol Biol 31, 1589–1600 (2024).

16. A. C. Carter, et al., Spen links RNA-mediated endogenous retrovirus silencing and X chromosome inactivation. Elife 9, e54508 (2020).

17. A. Monfort, et al., Identification of *Spen* as a Crucial Factor for *Xist* Function through Forward Genetic Screening in Haploid Embryonic Stem Cells. Cell Reports 12, 554–561 (2015).

18. F. Dossin, et al., SPEN integrates transcriptional and epigenetic control of X-inactivation. Nature 578, 455–460 (2020).

19. A. Hauth, et al., Escape from X inactivation is directly modulated by Xist noncoding RNA. Nat Cell Biol 28, 166–181 (2026).

20. A. C. Button, S. D. Hall, E. L. Ashley, C. A. McHugh, Dissection of protein and RNA regions required for SPEN binding to XIST A-repeat RNA. RNA 30, 240–255 (2024).

21. S. Collombet, et al., RNA polymerase II depletion from the inactive X chromosome territory is not mediated by physical compartmentalization. Nat Struct Mol Biol 30, 1216–1223 (2023).

22. J. W. Jachowicz, et al., Xist spatially amplifies SHARP/SPEN recruitment to balance chromosome-wide silencing and specificity to the X chromosome. Nat Struct Mol Biol 29, 239–249 (2022).

23. T. B. Nesterova, et al., Systematic allelic analysis defines the interplay of key pathways in X chromosome inactivation. Nat Commun 10, 3129 (2019).

24. J. B. Trotman, et al., Xist Repeat A coordinates an assembly of SR proteins to recruit SPEN and induce gene silencing. bioRxiv 2025.05.21.655143 (2025). 10.1101/2025.05.21.655143.

25. A. Pandya-Jones, et al., A protein assembly mediates Xist localization and gene silencing. Nature 587, 145–151 (2020).

26. J. S. Bowness, et al., Xist-mediated silencing requires additive functions of SPEN and Polycomb together with differentiation-dependent recruitment of SmcHD1. Cell Rep 39, 110830 (2022).

27. R. Weigert, et al., Dynamic antagonism between key repressive pathways maintains the placental epigenome. Nat Cell Biol 25, 579–591 (2023).

28. O. Masui, et al., Polycomb repressive complexes 1 and 2 are each essential for maintenance of X inactivation in extra-embryonic lineages. Nat Cell Biol 25, 134–144 (2023).

29. D. Andergassen, Z. D. Smith, H. Kretzmer, J. L. Rinn, A. Meissner, Diverse epigenetic mechanisms maintain parental imprints within the embryonic and extraembryonic lineages. Dev Cell 56, 2995–3005.e4 (2021).

30. M. Almeida, et al., PCGF3/5-PRC1 initiates Polycomb recruitment in X chromosome inactivation. Science 356, 1081–1084 (2017).

31. K. Kuroda, et al., Regulation of marginal zone B cell development by MINT, a suppressor of Notch/RBP-J signaling pathway. Immunity 18, 301–312 (2003).

32. J. J. Żylicz, et al., The Implication of Early Chromatin Changes in X Chromosome Inactivation. Cell 176, 182–197.e23 (2019).

33. A. Guyochin, et al., Live cell imaging of the nascent inactive X chromosome during the early differentiation process of naive ES cells towards epiblast stem cells. PLoS One 9, e116109 (2014).

34. H. Shiura, K. Abe, Xist/Tsix expression dynamics during mouse peri-implantation development revealed by whole-mount 3D RNA-FISH. Sci Rep 9, 3637 (2019).

35. V. Mutzel, et al., A symmetric toggle switch explains the onset of random X inactivation in different mammals. Nat Struct Mol Biol 26, 350–360 (2019).

36. G. Pacini, et al., Integrated analysis of Xist upregulation and X-chromosome inactivation with single-cell and single-allele resolution. Nat Commun 12, 3638 (2021).

37. I. Okamoto, et al., Eutherian mammals use diverse strategies to initiate X-chromosome inactivation during development. Nature 472, 370–374 (2011).

38. I. Okamoto, et al., The X chromosome dosage compensation program during the development of cynomolgus monkeys. Science 374, eabd8887 (2021).

39. A. Loda, S. Collombet, E. Heard, Gene regulation in time and space during X-chromosome inactivation. Nat Rev Mol Cell Biol 23, 231–249 (2022).

40. E. Kanata, R. Duffié, E. G. Schulz, Establishment and maintenance of random monoallelic expression. Development 151, dev201741 (2024).

41. R. M. Malcore, S. Kalantry, A Comparative Analysis of Mouse Imprinted and Random X-Chromosome Inactivation. Epigenomes 8, 8 (2024).

42. J. Starmer, T. Magnuson, A new model for random X chromosome inactivation. Development 136, 1–10 (2009).

43. C. Gontan, et al., RNF12 initiates X-chromosome inactivation by targeting REX1 for degradation. Nature 485, 386–390 (2012).

44. I. Jonkers, et al., RNF12 is an X-Encoded dose-dependent activator of X chromosome inactivation. Cell 139, 999–1011 (2009).

45. N. C. Rezende, et al., Rex1 (Zfp42) null mice show impaired testicular function, abnormal testis morphology, and aberrant gene expression. Dev Biol 356, 370–382 (2011).

46. J. Shin, et al., RLIM is dispensable for X-chromosome inactivation in the mouse embryonic epiblast. Nature 511, 86–89 (2014).

47. J. Lin, et al., KDM6A facilitates Xist upregulation at the onset of X inactivation. Biol Sex Differ 16, 1 (2025).

48. M. K. Samanta, et al., Activation of Xist by an evolutionarily conserved function of KDM5C demethylase. Nat Commun 13, 2602 (2022).

49. T. Schwämmle, et al., Reporter CRISPR screens decipher cis-regulatory and trans-regulatory principles at the Xist locus. Nat Struct Mol Biol 32, 2465–2475 (2025).

50. J. T. Lee, W. M. Strauss, J. A. Dausman, R. Jaenisch, A 450 kb transgene displays properties of the mammalian X-inactivation center. Cell 86, 83–94 (1996).

51. A. Wutz, T. P. Rasmussen, R. Jaenisch, Chromosomal silencing and localization are mediated by different domains of Xist RNA. Nat Genet 30, 167–174 (2002).

52. T. Robert-Finestra, et al., SPEN is required for Xist upregulation during initiation of X chromosome inactivation. Nat Commun 12, 7000 (2021).

53. J. Kim, S. Yoo, Software review: DEAP (Distributed Evolutionary Algorithm in Python) library. Genet Program Evolvable Mach 20, 139–142 (2019).

54. A. E. Pomeroy, et al., A predictive model of gene expression reveals the role of network motifs in the mating response of yeast. Science Signaling 14, eabb5235 (2021).

55. J. J. Tyson, K. C. Chen, B. Novak, Sniffers, buzzers, toggles and blinkers: dynamics of regulatory and signaling pathways in the cell. Curr Opin Cell Biol 15, 221–231 (2003).

56. M. Webster, K. L. Witkin, O. Cohen-Fix, Sizing up the nucleus: nuclear shape, size and nuclear-envelope assembly. J Cell Sci 122, 1477–1486 (2009).

57. G. Yahya, et al., Sublinear scaling of the cellular proteome with ploidy. Nat Commun 13, 6182 (2022).

58. E. J. Sousa, et al., Exit from Naive Pluripotency Induces a Transient X Chromosome Inactivation-like State in Males. Cell Stem Cell 22, 919–928.e6 (2018).

59. C. Vallot, et al., XACT Noncoding RNA Competes with XIST in the Control of X Chromosome Activity during Human Early Development. Cell Stem Cell 20, 102–111 (2017).

60. A. Mathur, R. Ghosh, A. Nunes-Alves, Recent Progress in Modeling and Simulation of Biomolecular Crowding and Condensation Inside Cells. J. Chem. Inf. Model. 64, 9063–9081 (2024).

61. K. L. Saar, et al., Theoretical and Data-Driven Approaches for Biomolecular Condensates. Chem Rev 123, 8988–9009 (2023).

62. A. Wutz, R. Jaenisch, A shift from reversible to irreversible X inactivation is triggered during ES cell differentiation. Mol Cell 5, 695–705 (2000).

63. J. M. Engreitz, et al., The Xist lncRNA Exploits Three-Dimensional Genome Architecture to Spread Across the X Chromosome. Science 341, 1237973 (2013).

64. Z. Du, et al., Stepwise de novo establishment of inactive X chromosome architecture in early development. Nat Genet 56, 2185–2198 (2024).

65. S. Gayen, E. Maclary, M. Hinten, S. Kalantry, Sex-specific silencing of X-linked genes by Xist RNA. Proceedings of the National Academy of Sciences 113, E309–E318 (2016).

66. S. Gayen, E. Maclary, E. Buttigieg, M. Hinten, S. Kalantry, A Primary Role for the Tsix lncRNA in Maintaining Random X-Chromosome Inactivation. Cell Rep 11, 1251–1265 (2015).

67. J. M. Kim, D. R. Larson, Timing is everything: transcription bursting in development. Genes Dev (2025). 10.1101/gad.352465.124.

68. E. A. Leyes Porello, R. T. Trudeau, B. Lim, Transcriptional bursting: stochasticity in deterministic development. Development 150, dev201546 (2023).

69. M. Nagel, D. J. Taatjes, Regulation of RNA polymerase II transcription through re-initiation and bursting. Mol Cell 85, 1907–1919 (2025).

70. J.-G. Chiou, et al., Principles that govern competition or co-existence in Rho-GTPase driven polarization. PLOS Computational Biology 14, e1006095 (2018).

71. A. D. Pourmorady, et al., RNA-mediated symmetry breaking enables singular olfactory receptor choice. Nature 625, 181–188 (2024).

72. D. U. Kissiov, et al., Binary outcomes of enhancer activity underlie stable random monoallelic expression. Elife 11, e74204 (2022).

73. S. N. Kravitz, et al., Random allelic expression in the adult human body. Cell Rep 42, 111945 (2023).

74. V. Tucci, A. R. Isles, G. Kelsey, A. C. Ferguson-Smith, Erice Imprinting Group, Genomic Imprinting and Physiological Processes in Mammals. Cell 176, 952–965 (2019).

75. M. E. Royce-Tolland, et al., The A-repeat links ASF/SF2-dependent Xist RNA processing with random choice during X inactivation. Nat Struct Mol Biol 17, 948–954 (2010).

76. C. An, et al., Overcoming Autocrine FGF Signaling-Induced Heterogeneity in Naive Human ESCs Enables Modeling of Random X Chromosome Inactivation. Cell Stem Cell 27, 482–497.e4 (2020).

77. T. W. Theunissen, et al., Molecular Criteria for Defining the Naive Human Pluripotent State. Cell Stem Cell 19, 502–515 (2016).

78. R. M. Karvas, et al., Stem-cell-derived trophoblast organoids model human placental development and susceptibility to emerging pathogens. Cell Stem Cell 29, 810–825.e8 (2022).

79. S. A. Khan, T. W. Theunissen, Modeling X-chromosome inactivation and reactivation during human development. Curr Opin Genet Dev 82, 102096 (2023).

80. G. Furlan, R. Galupa, Mechanisms of Choice in X-Chromosome Inactivation. Cells 11, 535 (2022).

81. F.-A. Fortin, F.-M. De Rainville, M.-A. G. Gardner, M. Parizeau, C. Gagné, DEAP: evolutionary algorithms made easy. J. Mach. Learn. Res. 13, 2171–2175 (2012).

82. J. C. Herron, et al., Spatial models of pattern formation during phagocytosis. PLOS Computational Biology 18, e1010092 (2022).

83. S. S. Andrews, D. Bray, Stochastic simulation of chemical reactions with spatial resolution and single molecule detail. Phys Biol 1, 137–151 (2004).

84. N. Mukherjee, et al., Integrative classification of human coding and noncoding genes through RNA metabolism profiles. Nat Struct Mol Biol 24, 86–96 (2017).

85. G. Wei, et al., m6A and the NEXT complex direct Xist RNA turnover and X-inactivation dynamics. Nat Struct Mol Biol 32, 2242–2251 (2025).

86. S. A. Sheardown, et al., Stabilization of Xist RNA mediates initiation of X chromosome inactivation. Cell 91, 99–107 (1997).

87. M. Y. Hein, et al., A human interactome in three quantitative dimensions organized by stoichiometries and abundances. Cell 163, 712–723 (2015).

88. X. Zhang, et al., An Interaction Landscape of Ubiquitin Signaling. Mol Cell 65, 941–955.e8 (2017).

89. J. R. Wiśniewski, M. Y. Hein, J. Cox, M. Mann, A “proteomic ruler” for protein copy number and concentration estimation without spike-in standards. Mol Cell Proteomics 13, 3497–3506 (2014).

90. A. J. Brenes, et al., Proteomic and functional comparison between human induced and embryonic stem cells. eLife (2024). Available at: https://elifesciences.org/articles/92025/figures [Accessed 19 February 2026].

91. D. Smeets, et al., Three-dimensional super-resolution microscopy of the inactive X chromosome territory reveals a collapse of its active nuclear compartment harboring distinct Xist RNA foci. Epigenetics Chromatin 7, 8 (2014).

