## Supplemental Material for "A three-factor model can explain symmetry breaking and the n-1 rule during X-chromosome inactivation"

**This PDF file includes:**

Supporting text
Figures S1 to S7
Tables S1 to S2
Legends for Movies S1 to S3

**Other supporting materials for this manuscript include the following:**

Movies S1 to S3

### Supporting Information Text

#### Evolutionary algorithm for parameter optimization

To optimize the parameters in our model, we adapted the evolutionary algorithm (EA) from Pomeroy et al. that uses DEAP (Distributed Evolutionary Algorithms in Python) (1–3). EA is a biologically inspired optimization method that iteratively improves candidate parameter sets based on principles of natural selection. The algorithm begins by initializing a population of parameter sets, sampled from biologically realistic ranges (Table 1 and Supporting Information). Each parameter set is evaluated by simulating the ODE model and using the simulation results to compute an appropriately constructed fitness score (Supporting Information). Parameters sets are then updated using mutation and recombination to generate better solutions over multiple generations.

#### Parameter bounds

Based on the results of several studies, the total number of *Xist* molecules per chromosome has been estimated to be on the order of 100 molecules (4–8). Therefore, we allowed the number of *Xist* binding sites ($X_{bs}$) to range between 100-200. Values for $X_{bs}$ found by the EA for both models were all close to 100.

SPEN expression levels exhibit high variability across different cell types and high cell-to-cell variability within specific cell types. Quantitative proteomic studies report SPEN copy numbers ranging from 35–170 in mouse embryonic stem cells (9) to ~4,600 in HeLa cells (10), and up to 15,000 in certain human cancer lines (11). Recent data from human cells show an average of 4182 for human embryonic stem cells and 6,215 copies for human induced pluripotent stem cells (hiPSCs)) (12). To accommodate this large variability, our range for SPEN abundance ($s_T$) was set broadly from $10^{1.5}$ to $10^{4.5}$ (~30-30,000). The median value of $s_T$ returned by the EA was approximately 3000 (ranging from 50-30000), which is consistent with values reported in the literature. Markaki et al. (2021) demonstrated that individual *Xist*-associated supermacromolar complexes contain roughly 2 *Xist* RNAs and up to 35 SPEN molecules (5). In the EA we let the number of SPEN that can associate with a single *XIST* molecule ($N_s$) equal to round($s_T$/ $X_{bs}$). The values found by the EA ranged between 1 and 50 molecules, which is consistent with this observed stoichiometry.

We allowed the degradation rate of free *Xist* ($\delta_x$) to range between $10^{-2.5}$ -$10^{-1.5}$ $min^{-1}$ (~0.003-0.03 $min^{-1}$). This range is from multiple independent, time-resolved transcriptomic and transcriptional inhibition studies across different cell lines. Mukherjee et al. (2017) determined the degradation rate of *XIST* in human HEK293 to be in the range 0.006 - 0.01 $min^{-1}$ (13). Similarly, single-cell transcriptomics by Wei et al. (2025) and Rodermund et al. (2021) established that *Xist* RNA turnover in mouse embryonic stem cells (ESCs) occurs within 140 - 220 min, corresponding to a rate of roughly 0.004-0.007 $min^{-1}$ (6, 14). In early undifferentiated ESCs where *Xist* is not yet stably anchored, transcriptional inhibition reveals a rapid half-life of 20-45 minutes, translating to a degradation rate of 0.02-0.03 $min^{-1}$ (15, 16). Our chosen range for $\delta_x$ captures these experimental measurements. Mukherjee et al. (2017) measured the *Xist* synthesis rate to be around 1-10 TPM/min and *Xist* RNA processing rate to be 0.1-1 $min^{-1}$. Our models combine synthesis and processing rates together as one step. Therefore, the maximum *Xist* production rate ($\alpha_x$) was bounded between 0.001 and 10 $min^{-1}$.

For reaction rates related to the activator ($\alpha_A$, $\delta_A$, $K_A$), we used broad parameter ranges ($\alpha_A$ = [$10^{-3}$,$10^2$] $min^{-1}$, $\delta A$ = [$10^{-3}$,10] $min^{-1}$, $K_A$ = [1,$10^4$]) because the molecular identity of this factor is unknown. However, these kinetic parameters are constrained because they must produce activator dynamics that occurs on time scales consistent with that of *XIST* and SPEN and lead to XCI on biologically relevant time scales. The parameter ranges found by the EA can serve as kinetic clues to guide the experimental identification of the activator.

An assumption of the model is that SPEN stabilizes *Xist* on X-chromosomes. This effect is captured in the model by the *XIST* dissociation rate being a decreasing function of bound SPEN. Using the steady state values of bound SPEN in this function, the median dissociation rate for all the parameter sets identified by the EA was 0.05 $min^{-1}$. This value is consistent with results based on FRAP measurements (0.052 $min^{-1}$) reported by Markaki et. al (2021). For the rate constant $k_4$ for SPEN dissociating from *Xist*, Markaki et al. (2021) reported biphasic dissociation kinetics for SPEN from *Xist*, identifying distinct fast (40 $min^{-1}$) and slow (0.4 $min^{-1}$) exchanging populations. We used a range for $k_4$ between 0.1 – $10^{1.6}$ $min^{-1}$. The EA returned values for $k_4$ that spanned this range.

In-vivo association rates for ribonucleoprotein complexes are difficult to measure. Association rates for *Xist* with chromatin ($k_1$) and SPEN with *Xist* ($k_3$) are governed by local molecular crowding, chromatin conformational change and phase separation (5, 6, 17). Because these local dynamics cause effective association rates that span orders of magnitude, we provided the EA with a broad range of values ($10^{-4}$ to 10 $min^{-1}$), capturing reaction-limited binding to rapid diffusion-limited binding. Because the opposing dissociation rates ($k_2$, $k_4$) are anchored to measured in vivo lifetimes, the association rates found by the EA must produce time scales for XCI to occur on biologically realistic time scales.

The model contains 3 Hill functions for: 1) the induction of *Xist* by the activator, 2) the stabilization of *Xist* on X-chromosomes by SPEN and 3) the repression of activator expression by SPEN. For simplicity, we took the Hill coefficient in the term for the induction of *XIST* by the activator to be 1. We bound the other two Hill coefficients (*m* and *n*) between 1 (hyperbolic or Michaelis-Menten kinetics) and 10 (switch-like kinetics). Half-maximal concentrations ($K_s$, $K_A$, $K_n$) were bounded from 1 to 10000 molecules, ensuring the optimization searched exclusively within the physiological intracellular dynamic range defined by global proteomics (9–12).

**Fitness scores used in the evolutionary algorithm**

For the male diploid cell (single-X case), the fitness score is defined as:

$$Score_1 = X_{b1} + Penalty_1,$$

where $X_{b1}$ is the steady-state number of bound *Xist* on the single X chromosome, which should be minimized to prevent XCI. The penalty term $Penalty_1 = 100000 \times \left|X_{b1,\ t=10000min} - X_{b1,\ t=6000min}\right|$ ensures that that bound *Xist* reaches a steady state within 6000 minutes, in alignment with prior studies indicating *Xist* clusters stabilize within one to five days in mouse embryonic stem cells (5, 6, 18, 19). The initial bound *Xist* is set equal to the total number of *Xist* binding sites on one chromosome ($X_{bs}$), but at steady-state the amount of bound *Xist* should be minimal.

For female diploid cells (XX), the fitness score is defined as:

$$Score_2 = \frac{X_{bs}}{1+|X_{b2}-X_{b1}|} + Penalty_2,$$

where $X_{b2} - X_{b1}$ represents the difference in the steady-state number of bound *Xist* between the two chromosomes, which should be maximized to ensure robust XCI asymmetry. Similarly, the penalty term is given by: $Penalty_2 = 100000 \times \sum_{i=1}^{2}\left|X_{bi,\ t=10000min} - X_{bi,\ t=6000min}\right|$. We used two initial conditions: $X_{b2} = 2$, $X_{b1} = 1$; and $X_{b2} = X_{bs}$, $X_{b1} = X_{bs} - 1$, with their scores summed. This ensures that a chromosome with a slight initial advantage in bound *Xist* accumulates significantly more *Xist* than the other at steady state.

For female diploid cells with three X chromosomes (XXX), the fitness function remains the same as for the XX case, but the penalty term is extended to include the third chromosome: $Penalty_3 = 100000 \times \sum_{i=1}^{3}\left|X_{bi,\ t=10000min} - X_{bi,\ t=6000min}\right|$. We applied three initial conditions: $X_{b3} = 3,\ X_{b2} = 2, X_{b1} = 1$; $X_{b3} = X_{bs}, X_{b2} = X_{bs} - 1$, $X_{b1} = X_{bs} - 2$; and $X_{b3} = X_{bs}, X_{b2} = 2, X_{b1} = 1$. The two chromosomes with more initially bound *Xist* ($X_{b2}$ and $X_{b3}$) should accumulate *Xist* while avoiding dominance by a single chromosome ($X_{b3}$).

The overall fitness is given by the sum of the three score functions:

$$Final\ score = Score_1 + Score_2 + Score_3.$$

A threshold fitness score of 50 was set as the criterion for a good solution. The EA successfully identified parameter sets generating the required behavior (Table 1).

To search parameter space for the SPEN depletion model, we added a penalty term for free SPEN:

$$Penalty_{free\ SPEN} = 0.1\ (S_T - \sum_{j=1}^{N_c} S_{bj}).$$

For the activation inhibition model, a penalty term for bound SPEN was added to ensure the mechanism of SPEN depletion did not contribute to XCI:

$$Penalty_{bound\ SPEN} = 0.1 \sum_{j=1}^{N_c} S_{bj}.$$

To test if the models can produce transient expression of *Xist* in male cells, we added the following term to the fitness score:

$$Score_{transient\ signal} = \frac{50}{1+X_{b1,max}-X_{b1}}.$$

where $X_{b1}$ is the steady-state number of bound *Xist* on the single X chromosome. $X_{b1,max}$ is the maximum number of bound *Xist*. The difference between $X_{b1,max}$ and $X_{b1}$ is maximized to allow the occurrence of transient *Xist* expression. The parameters values used to generate the results in Fig. 4 are given in Table 1.

**Particle based simulations**

The simulation domain representing the full nucleus was a cube measuring 16 µm on each side with periodic boundary conditions. Rate constants *k* for first-order reactions are the same as in the ODE model such that those reactions have a probability

$$P = 1 - \exp(-k\ \delta t) \approx k\ \delta t,$$

for sufficiently small $\delta t$. For second-order reactions, whenever two reactant particles come within a separation of ρ, the probability of reaction during a time interval $\delta t$ is

$$P = 1 - \exp(-\lambda\ \delta t) \approx \lambda\ \delta t,$$

for sufficiently small $\delta t$. We did not attempt to directly transform macroscopic rate constants into microscopic rates. Instead, we estimated the rates and tuned them by hand. The values for the rates used in the particle-based simulations are given in Table 2.

We assumed X chromosomes occupy spherical domains of radius around 2.5 µm. Within each sphere, 100 *Xist* binding sites were placed sequentially using a 3D random-walk with fixed step length of 1 µm. Starting from an initial seed position, each subsequent position ($x_i$, $y_i$, $z_i$) was generated by sampling spherical coordinates:

$$\theta \sim Uniform(0, 2\pi),\ \ \phi \sim Uniform(0, \pi),$$

and setting

$$x_i = x_{i-1} + \Delta \sin\phi \ \cos\theta,$$
$$y_i = y_{i-1} + \Delta \sin\phi \ \sin\theta,$$
$$z_i = z_{i-1} + \Delta \cos\phi,$$

where Δ = 1 μm. If the proposed point lay outside the sphere – specifically, if its distance from the chromosome center exceeded 2.49 μm – it was rejected and resampled. This procedure continued until 100 valid binding-site positions were obtained.

For the results presented in Fig. 5, we counted the total number of bound SPEN $S_{bj}$ on each chromosome every 0.1 min and calculated the corresponding dissociation rate for every *Xist*-SPEN complex on that chromosome. Each bound *Xist* can associate with a maximum number of 10 SPEN.

The number of activators $N_A$ inserted into the nucleus at each time step is:

$$N_A = dt \times \alpha_A \sum_{j=1}^{N_C} \frac{1}{\left(1+\left(\frac{S_{bj}}{K_n}\right)^n\right)}.$$

For each X chromosome the average *Xist* synthesis rate $s_{Xist}$ was calculated as:

$$s_{Xist} = \frac{\alpha_x A_x}{K_A + A_x}.$$

**Model using a common pool for *Xist***

The ODE model in the main text assumes *Xist* molecules diffuse slowly such that they only bind to the chromosome from which they are transcribed. To test if this assumption is required for the model to function correctly, we revised the equations assuming a common pool of *Xist*. This was accomplished by making the following modifications to Eqs. (2) and (3):

$$\frac{dX_f}{dt} = \frac{\alpha_x A_x}{K_A + A_x} - \delta_x X_f - k_1 \sum_{i=1}^{N_S} (X_{bs} - X_{bi}) x_f + k_2 \sum_{i=1}^{N_S} \frac{X_{bi}}{1 + \left(\frac{S_{bi}}{K_s}\right)^m}$$

$$\frac{dX_{bi}}{dt} = k_1 (X_{bs} - X_{bi}) x_f - k_2 \frac{X_{bi}}{1 + \left(\frac{S_{bi}}{K_s}\right)^m}$$

Simulations using these equations yielded similar parameter distributions and results for both mechanisms (Figs. S6 and S7 and Table S1). These results further demonstrate that a cis-regulatory network is not required for random XCI.

**Model with an alternative form of positive feedback**

The model presented in the main text relies on positive feedback through SPEN's stabilization of *Xist* on X chromosomes to amplify asymmetric accumulation of *Xist* molecules. We tested the ODE model with another form of SPEN-mediated positive feedback, in which SPEN enhances *Xist*-chromosome binding (Fig. S3A). This required modifying Eqs. (2) and (3) in the following way:

$$\frac{dX_{fi}}{dt} = \frac{\alpha_x A_x}{K_A + A_x} - \delta_x X_{fi} - (X_{bs} - X_{bi}) x_{fi} (k_1 + k_5 \frac{S_{bi}^m}{S_{bi}^m + K_S^m}) + k_2 X_{bi}$$

$$\frac{dX_{bi}}{dt} = (X_{bs} - X_{bi}) x_{fi} (k_1 + k_5 \frac{S_{bi}^m}{S_{bi}^m + K_S^m}) - k_2 X_{bi}$$

where $k_1$ is the basal rate of *Xist* binding to the chromosome and $k_5 \frac{S_{bi}^m}{S_{bi}^m + K_S^m}$ denotes SPEN-mediated positive feedback of *Xist*-chromosome binding. This form of positive feedback produced similar results as the original model (Fig. S3, activator inhibition and Fig. S4, SPEN depletion). Note that for SPEN depletion to play a dominant role, the rate of positive feedback ($k_5$) in the model must be significantly higher than that in the activator inhibition model, which is demonstrated by Fig. S4D. These alternative models provide support for the robustness of the combined positive and negative feedback regulatory network for XCI.

### Supplemental Figures

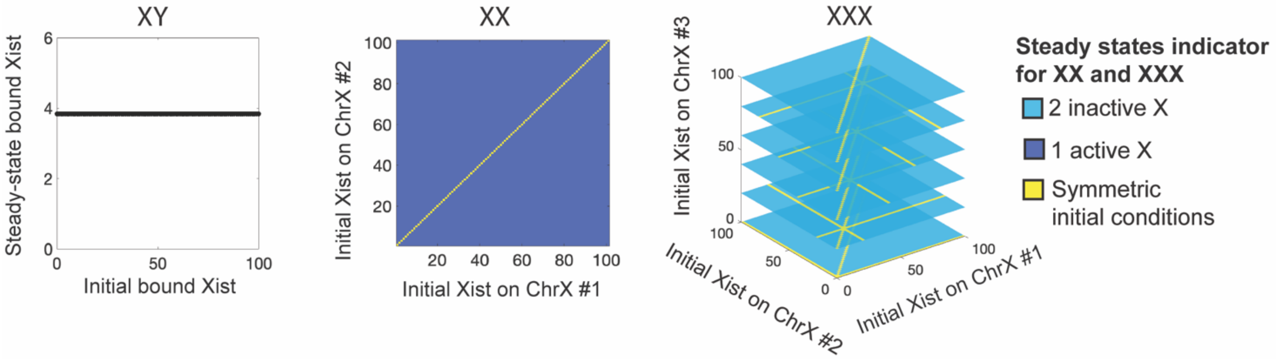


**Figure S1**. Plots demonstrating that the n-1 rule is satisfied independent of initial conditions for the cases of XY, XX and XXX for the activator inhibition model.

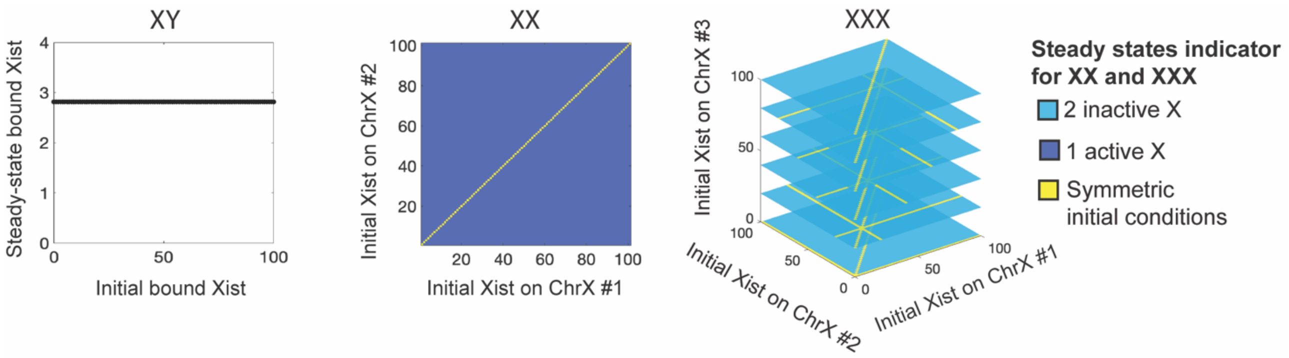


**Figure S2**. Plots demonstrating that the n-1 rule is satisfied independent of initial conditions for the cases of XY, XX and XXX for the SPEN depletion model.

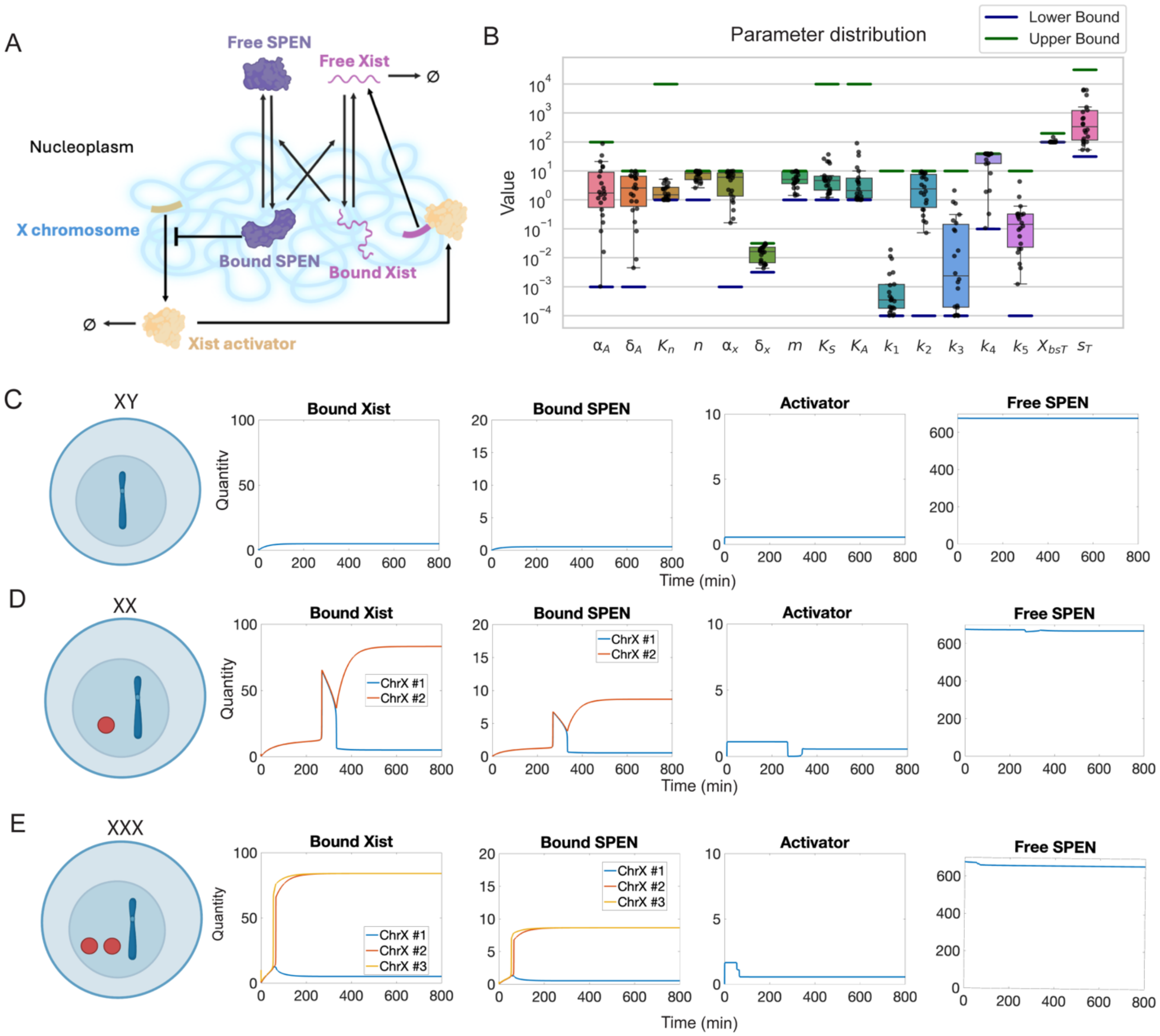


**Figure S3**. Simulation results for the activator inhibition model with mutual activation. **A)** Model schematic. Created with BioRender.com. Same as Fig. 1A except that SPEN accelerate *Xist*-chromosome binding instead of inhibiting *Xist*-chromosome dissociation. **B**) Parameter sets identified by evolutionary algorithm. See Table S2 for parameter descriptions and relevant references. **C)** Time series for number of molecules of bound *Xist*, bound SPEN, activator and free SPEN for the case of XY males. **D)** Same as **C** except for XX females. **E)** Same as **C** except for XXX females.

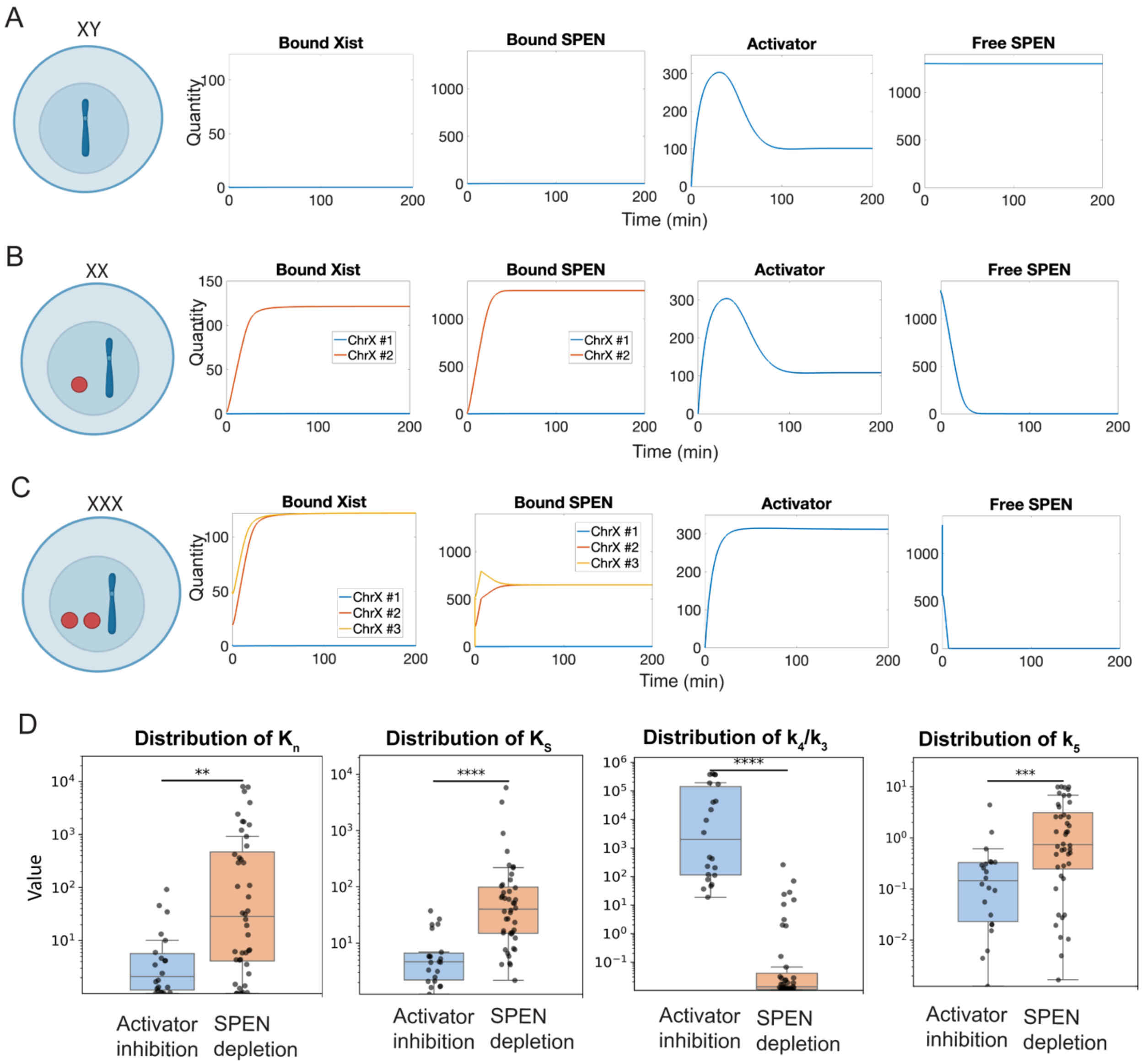


**Figure S4.** Results for SPEN depletion model with mutual activation. **A**) Time series for number of molecules of bound *Xist*, bound SPEN, activator and free SPEN for XX females. **B**) Same as **A** except for the case of XY males. **C**) Same as **A** except for XXX females. **D**) Distributions of parameters that are significantly different between the SPEN depletion and activator inhibition models (* p ≤ 0.05, ** p ≤ 0.01, *** p ≤ 0.001, **** p ≤ 0.0001). Y axes are in $\log_{10}$ scale.

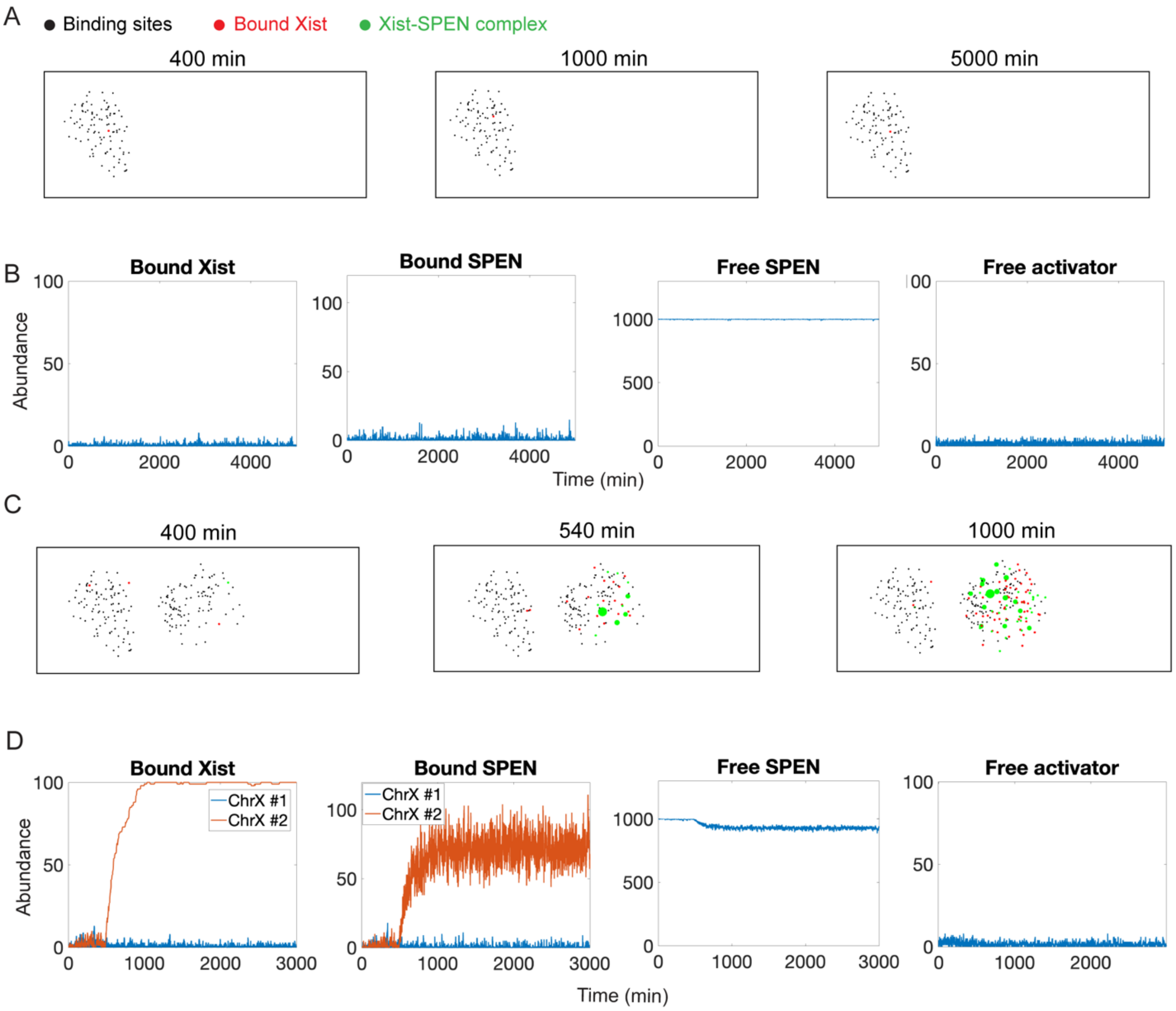


**Figure S5**. Particle-based simulation results for the case of single X-chromosome (**A** and **B**) and two X-chromosomes (**C** and **D**). **A**) and **C**) Spatial temporal distribution of *Xist* binding sites, molecules of bound *Xist*, and molecules of bound SPEN. **B**) and **D**) Time series for the molecular abundance of bound *Xist*, bound SPEN, free SPEN, and activators.

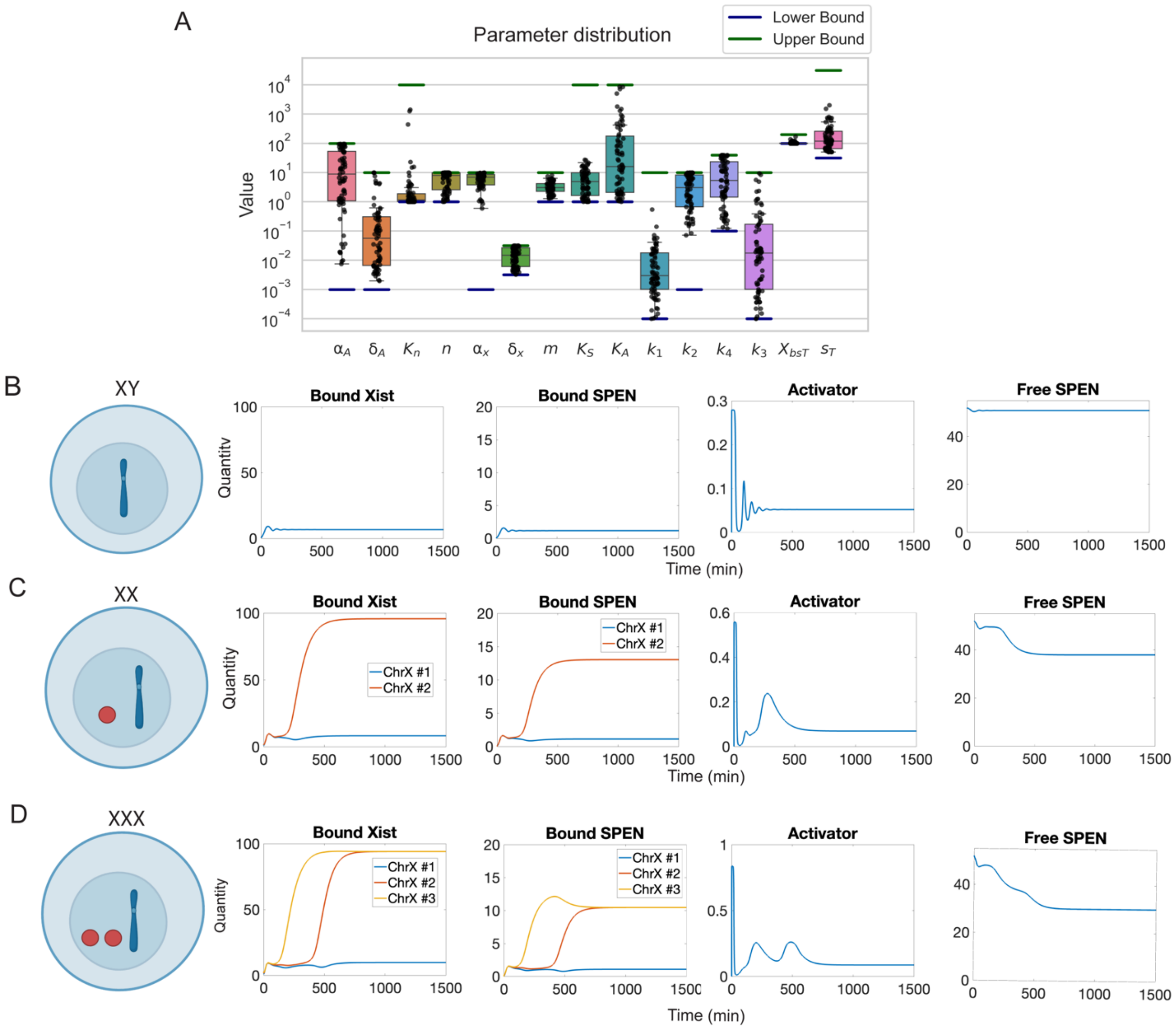


**Figure S6**. Simulation results for the activation inhibition model with a common pool of *Xist* molecules. **A)** Parameter sets identified by evolutionary algorithm. **B)** Time series for number of molecules of bound *Xist*, bound SPEN, activator and free SPEN for the case of XY males. **C)** Same as **B** except for XX females. **D)** Same as **B** except for XXX females.

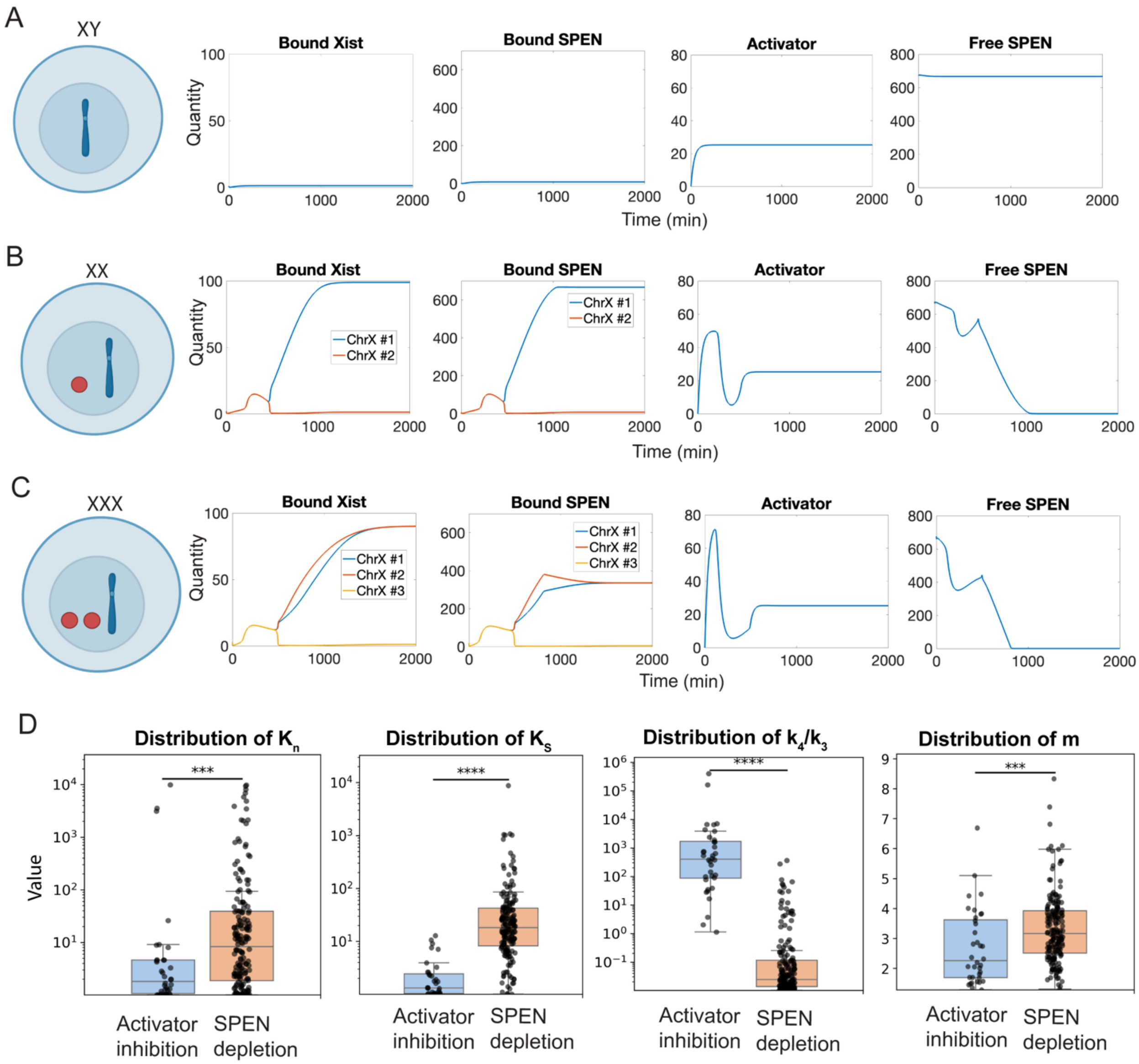


**Figure S7**. SPEN depletion model with a common pool of *Xist* molecules. **A**) Time series for number of bound *Xist*, bound SPEN, activator and free SPEN for XX females. **B**) Same as **A** except for the case of XY males. **C**) Same as **A** except for XXX females. **D**) Comparison of distributions of parameters between the models (* $p \leq 0.05$, ** $p \leq 0.01$, *** $p \leq 0.001$, **** $p \leq 0.0001$). Y axes of the first three panels are in $\log_{10}$ scale. Y axis of the last panel is in linear scale.

### Supplemental Tables

**Table S1**. Parameters for the model with a common pool of *Xist* molecules.

| Parameter description | Symbol | Search range | Parameter values used in Fig. S6 | Parameter values used in Fig. S7 |
|---|---|---|---|---|
| *Xist*-chromosome association rate | $k_1$ | $[10^{-4}, 10]$ min$^{-1}$ | 0.00084 min$^{-1}$ | 0.0018 min$^{-1}$ |
| *Xist*-chromosome dissociation rate in the absence of SPEN | $k_2$ | $[10^{-3},10]$ min$^{-1}$ | 0.27 min$^{-1}$ | 1.42 min$^{-1}$ |
| SPEN-*Xist* association rate | $k_3$ | $[10^{-4},10]$ min$^{-1}$ | 0.0015 min$^{-1}$ | 9.9 min$^{-1}$ |
| SPEN-*Xist* dissociation rate | $k_4$ | $[10^{-1},10^{1.6}]$ min$^{-1}$ | 0.36 min$^{-1}$ | 0.10 min$^{-1}$ |
| Maximum *Xist* transcription rate | $\alpha_x$ | $[10^{-3},10]$ min$^{-1}$ | 7.4 min$^{-1}$ | 3.9 min$^{-1}$ |
| Degradation rate for free *Xist* | $\delta_x$ | $[10^{-2.5},10^{-1.5}]$ min$^{-1}$ | 0.014 min$^{-1}$ | 0.027 min$^{-1}$ |
| Maximum activator production rate | $\alpha_A$ | $[10^{-3},10^{2}]$ min$^{-1}$ | 1.2 min$^{-1}$ | 0.707 min$^{-1}$ |
| Degradation rate for the activator | $\delta_A$ | $[10^{-3},10]$ min$^{-1}$ | 4.3 min$^{-1}$ | 0.028 min$^{-1}$ |
| Number of activator molecules at which the *Xist* transcription rate is half max | $K_A$ | $[1,10^{4}]$ | 2.19 molecules | 351 molecules |
| Number of SPEN molecules at which the *Xist* degradation rate is half max | $K_S$ | $[1,10^{4}]$ | 1.23 molecules | 26.8 molecules |
| Hill coefficient for SPEN inhibition of *Xist* dissociation | m | [1,10] | 2.58 | 2.58 |
| Number of SPEN molecules at which the activator production rate is half max | $K_n$ | $[1,10^{4}]$ | 1.0 molecules | 57.8 molecules |
| Hill coefficient for SPEN inhibition of activator production | n | [1,10] | 10.0 | 4.39 |
| Total *Xist* binding sites per chromosome | $X_{bs}$ | $[10^{2},10^{2.3}]$ | 100 molecules | 101 molecules |
| Total SPEN abundance | $S_T$ | $[10^{1.5},10^{4.5}]$ | 52 molecules | 676 molecules |

| | | | | |
|---|---|---|---|---|
| Maximum number of SPEN that can bind to an *Xist* molecule | $N_S$ | $round(\frac{S_T}{X_{bs}})$ | 1 molecule | 7 molecules |

**Table S2**. Parameters for the model with mutual activation.

| Parameter description | Symbol | Search range | Parameter values used in Fig. S3 | Parameter values used in Fig. S4 |
|---|---|---|---|---|
| *Xist*-chromosome association rate | $k_1$ | $[10^{-4}, 10]$ $min^{-1}$ | 0.011 $min^{-1}$ | 0.0001 $min^{-1}$ |
| *Xist*-chromosome dissociation rate | $k_2$ | $[10^{-3},10]$ $min^{-1}$ | 8.57 $min^{-1}$ | 9.28 $min^{-1}$ |
| SPEN-*Xist* association rate | $k_3$ | $[10^{-4},10]$ $min^{-1}$ | 0.00019 $min^{-1}$ | 5.91 $min^{-1}$ |
| SPEN-*Xist* dissociation rate | $k_4$ | $[10^{-1},10^{1.6}]$ $min^{-1}$ | 8.39 $min^{-1}$ | 0.11 $min^{-1}$ |
| Rate for SPEN-enhanced *Xist*–chromosome binding | $k_5$ | $[10^{-4},10^{1}]$ $min^{-1}$ | 1.28 $min^{-1}$ | 2.57 $min^{-1}$ |
| Maximum *Xist* transcription rate | $\alpha_x$ | $[10^{-3},10]$ $min^{-1}$ | 10 $min^{-1}$ | 6.01 $min^{-1}$ |
| Degradation rate for free *Xist* | $\delta_x$ | $[10^{-2.5},10^{-1.5}]$ $min^{-1}$ | 0.029 $min^{-1}$ | 0.028 $min^{-1}$ |
| Maximum activator production rate | $\alpha_A$ | $[10^{-3},10^{2}]$ $min^{-1}$ | 1.89 $min^{-1}$ | 37.8 $min^{-1}$ |
| Degradation rate for the activator | $\delta_A$ | $[10^{-3},10]$ $min^{-1}$ | 3.43 $min^{-1}$ | 0.12 $min^{-1}$ |
| Number of activator molecules at which the *Xist* transcription rate is half max | $K_A$ | $[1,10^{4}]$ | 4.15 molecules | 31.1 molecules |
| Number of SPEN molecules at which the *Xist*-chromosome binding rate is half max | $K_S$ | $[1,10^{4}]$ | 5.78 molecules | 19.9 molecules |
| Hill coefficient for SPEN inhibition of *Xist* dissociation | m | [1,10] | 3.6 | 9.6 |
| Number of SPEN molecules at which the activator production rate is half max | $K_n$ | $[1,10^{4}]$ | 2.85 molecules | 2.23 molecules |

| | | | | |
|---|---|---|---|---|
| Hill coefficient for SPEN inhibition of activator production | n | [1,10] | 7.8 | 9.8 |
| Total *Xist* binding sites per chromosome | $X_{bs}$ | [$10^2$,$10^{2.3}$] | 101 molecules | 124 molecules |
| Total SPEN abundance | $S_T$ | [$10^{1.5}$,$10^{4.5}$] | 676 molecules | 1303 molecules |
| Maximum number of SPEN that can bind to an *Xist* molecule | $N_S$ | $round(\frac{S_T}{X_{bs}})$ | 7 molecules | 11 molecules |

**Movie S1**. Distribution of bound *Xist* (red), *Xist*-SPEN complexes (green), and *Xist* binding sites on the chromosomes (black) for the triple X case.

**Movie S2**. Distribution of bound *Xist* (red), *Xist*-SPEN complexes (green), and *Xist* binding sites on the chromosomes (black) for the female double-X case.

**Movie S3**. Distribution of bound *Xist* (red), *Xist*-SPEN complexes (green), and *Xist* binding sites on the chromosomes (black) for the male single-X case.

### References

1. F.-A. Fortin, F.-M. De Rainville, M.-A. G. Gardner, M. Parizeau, C. Gagné, DEAP: evolutionary algorithms made easy. *J. Mach. Learn. Res.* **13**, 2171–2175 (2012).

2. J. C. Herron, *et al.*, Spatial models of pattern formation during phagocytosis. *PLOS Computational Biology* **18**, e1010092 (2022).

3. A. E. Pomeroy, *et al.*, A predictive model of gene expression reveals the role of network motifs in the mating response of yeast. *Science Signaling* **14**, eabb5235 (2021).

4. A. Cerase, *et al.*, Spatial separation of Xist RNA and polycomb proteins revealed by superresolution microscopy. *Proceedings of the National Academy of Sciences* **111**, 2235–2240 (2014).

5. Y. Markaki, *et al.*, Xist nucleates local protein gradients to propagate silencing across the X chromosome. *Cell* **184**, 6212 (2021).

6. L. Rodermund, *et al.*, Time-resolved structured illumination microscopy reveals key principles of Xist RNA spreading. *Science* **372**, eabe7500 (2021).

7. D. Smeets, *et al.*, Three-dimensional super-resolution microscopy of the inactive X chromosome territory reveals a collapse of its active nuclear compartment harboring distinct Xist RNA foci. *Epigenetics Chromatin* **7**, 8 (2014).

8. H. Sunwoo, J. Y. Wu, J. T. Lee, The Xist RNA-PRC2 complex at 20-nm resolution reveals a low Xist stoichiometry and suggests a hit-and-run mechanism in mouse cells. *Proceedings of the National Academy of Sciences* **112**, E4216–E4225 (2015).

9. X. Zhang, *et al.*, An Interaction Landscape of Ubiquitin Signaling. *Mol Cell* **65**, 941-955.e8 (2017).

10. M. Y. Hein, *et al.*, A human interactome in three quantitative dimensions organized by stoichiometries and abundances. *Cell* **163**, 712–723 (2015).

11. J. R. Wiśniewski, M. Y. Hein, J. Cox, M. Mann, A “proteomic ruler” for protein copy number and concentration estimation without spike-in standards. *Mol Cell Proteomics* **13**, 3497–3506 (2014).

12. A. J. Brenes, *et al.*, Proteomic and functional comparison between human induced and embryonic stem cells. *eLife* (2024). Available at: https://elifesciences.org/articles/92025/figures [Accessed 19 February 2026].

13. N. Mukherjee, *et al.*, Integrative classification of human coding and noncoding genes through RNA metabolism profiles. *Nat Struct Mol Biol* **24**, 86–96 (2017).

14. G. Wei, *et al.*, m6A and the NEXT complex direct Xist RNA turnover and X-inactivation dynamics. *Nat Struct Mol Biol* **32**, 2242–2251 (2025).

15. S. A. Sheardown, *et al.*, Stabilization of Xist RNA mediates initiation of X chromosome inactivation. *Cell* **91**, 99–107 (1997).

16. A. Wutz, T. P. Rasmussen, R. Jaenisch, Chromosomal silencing and localization are mediated by different domains of Xist RNA. *Nat Genet* **30**, 167–174 (2002).

17. J. W. Jachowicz, *et al.*, Xist spatially amplifies SHARP/SPEN recruitment to balance chromosome-wide silencing and specificity to the X chromosome. *Nat Struct Mol Biol* **29**, 239–249 (2022).

18. K. Monkhorst, I. Jonkers, E. Rentmeester, F. Grosveld, J. Gribnau, X Inactivation Counting and Choice Is a Stochastic Process: Evidence for Involvement of an X-Linked Activator. *Cell* **132**, 410–421 (2008).

19. A. Wutz, R. Jaenisch, A shift from reversible to irreversible X inactivation is triggered during ES cell differentiation. *Mol Cell* **5**, 695–705 (2000).
